# Intralysosomal Amyloidogenesis and Proximity Labeling by Cathepsin C

**DOI:** 10.64898/2025.12.23.696283

**Authors:** Ruben D. Elias, Seth Allen, Idil I. Demiralp, Robert T. O’Neill, Jan-Hannes Schäfer, Hannah Siems, Elizabeth A. Montabana, Utz H. Ermel, Carl Ash, Fatema Abdurrob, Daniella A. Yacoubian, Oren L. Lederberg, Daniel Serwas, David A. Agard, Benjamin F. Cravatt, Jeffery W. Kelly

**Affiliations:** Department of Chemistry, Scripps Research; La Jolla, CA, USA; Department of Integrative Structural and Computational Biology Scripps Research; La Jolla, CA, USA; Biohub, San Francisco; Redwood City, CA, USA; Department of Biochemistry & Biophysics, University of California; San Francisco, CA, USA

## Abstract

The lysosome is a major catabolic organelle responsible for the breakdown of both intra- and extracellular substrates^1,2^. Lysosomal membrane damage mediated by pathologic amyloid fibrils is an area of recent focus^3-6^. The dipeptide ester LLOMe is typically employed to model lysosomal membrane damage^7-11^; however its mechanism of membranolysis was previously incompletely understood. Here, *in vitro* and cell-based analyses, and cryo-electron microscopy and tomography studies reveal LLOMe-derived oligopeptides generated by the lysosomal protease Cathepsin C assemble into cross-β-sheet amyloid fibrils within the lysosome. Additionally, we report lysosome membrane damage triggers the broadly nonspecific dipeptidyl ligase activity of Cathepsin C, facilitating the tagging of proximal proteins within the damaged lysosome lumen with a click chemistry handle: to our knowledge, the first reported localized proximity labeling approach exploiting a fully endogenous, non-engineered enzyme. While Cathepsin C ligase activity has been demonstrated *in vitro*^12,13^, our observations of dipeptidyl ligation onto proximal proteins in cells suggests an unexplored role of Cathepsin C in lysosomal biology and broadly exemplifies how other endogenous enzymes might be similarly exploited for proximity labeling. Altogether our results unveil two mechanisms by which dipeptide esters perturb lysosomal homeostasis and provide a roadmap for their utilization toward targeted studies of the lysosome.

---

The relationship between lysosome dysfunction and the onset and progression of neurodegeneration has become a major research focus^2,14-18^. A prion-like transmission of amyloid fibrils from affected to unaffected cells is proposed to drive pathology of certain amyloidoses including tauopathies and synucleopathies^19,20^, where endocytosis of extracellular amyloid fibril seeds, followed by endolysosomal breakout, mediates intracellular seeding^3,21-24^. Consequently, autolysosomal deacidification and lysosomal membrane damage are observed in murine and neuronal models of Alzheimer’s disease^4,5^. Endolysosomal amyloid fibril breakout is thought to be mediated by the membranolytic properties of amyloid assemblies^25^. This etiology is typically modeled by supplementing cultures with preformed amyloid fibrils, exposing cells for extended periods (1-2 days) before observing a given effect^6,21,23^, resulting in heterogeneity in the number of affected cells and in the subcellular localization of amyloid fibrils at a given timepoint.

L-leucyl-L-leucine methyl ester (LLOMe, Fig. 1a) has emerged as a routine biochemical reagent to induce rapid, acute lysosomal membrane damage upon exposure to cells, facilitating investigation of lysosomal repair mechanisms ^7-11,26-28^. The mechanism of action of LLOMe has previously been characterized to involve its ligation into (Leu-Leu)_n_-OH and (Leu-Leu)_n_-OMe oligopeptides (hereafter designated (Leu-Leu)_n_, where n=2-3 in lymphocytes^29^) catalyzed by the ligase activity of the lysosomal dipeptidase cathepsin C (CTSC), with emergent membranolytic properties^12,13,29-31^. Despite widespread use of LLOMe, the mechanistic basis of (Leu-Leu)_n_ membranolytic function has historically been unclear.

**Fig. 1.**
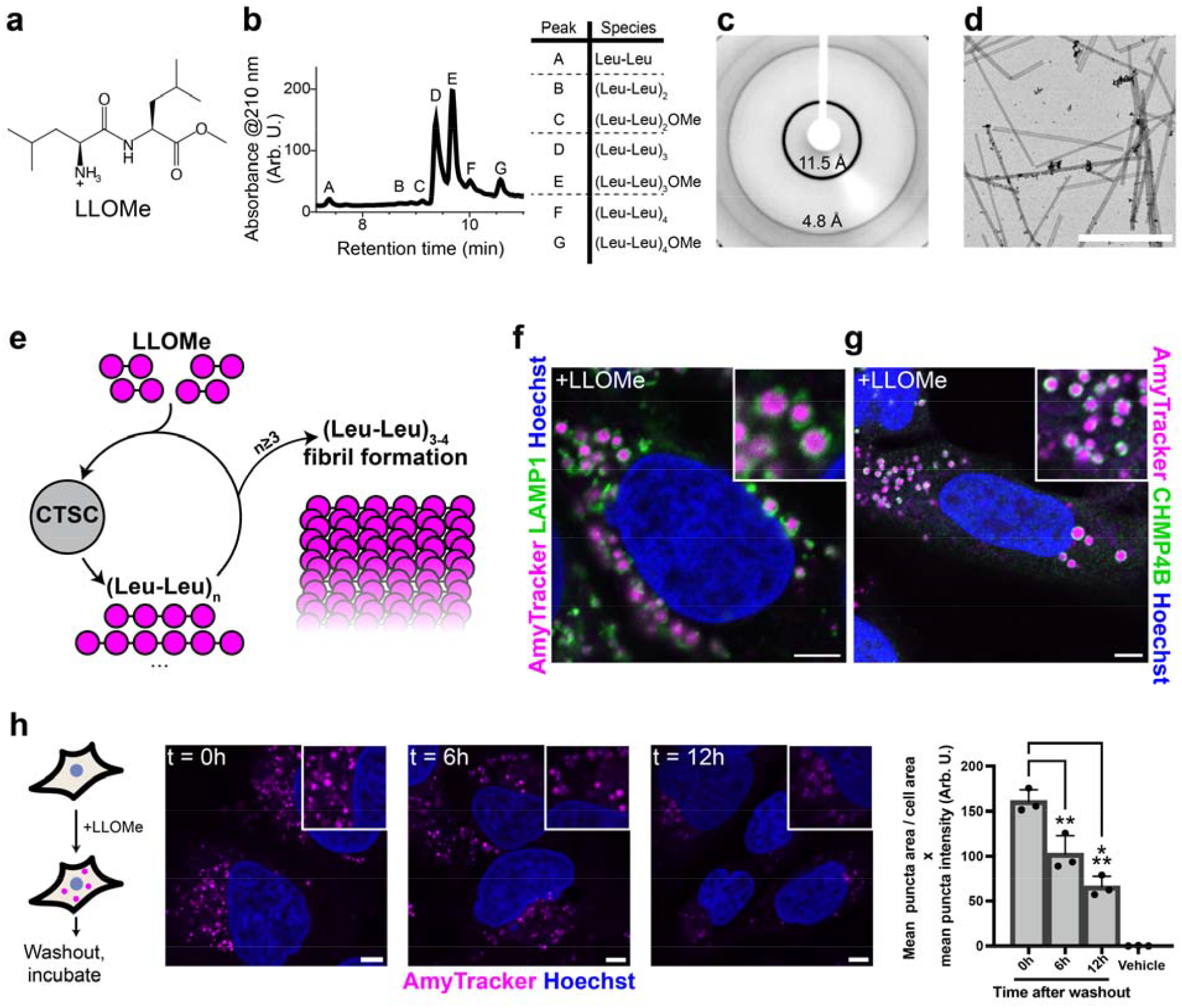
(Leu-Leu)_n_ oligopeptides spontaneously form cross β-sheet–rich amyloids within endolysosomes. **a**, Structure of LLOMe. **b**, LC-MS chromatogram of LLOMe (100 mM) + CTSC (500 nM) (in 20 mM sodium phosphate pH 6.5, 150 mM NaCl) overnight reaction precipitate dissolved in DMSO (left); assigned peaks identified by mass (right). **c**, X-ray diffraction pattern of LLOMe + CTSC reaction precipitate. **d**, Negative-stain transmission electron microscopy of LLOMe + CTSC reaction mixture. **e**, Model of CTSC-mediated LLOMe ligation and consequent fibril formation. **f**, Representative image of U-2 OS cells treated with LLOMe (1 mM, 10 minutes) fixed and stained against endolysosomal marker LAMP1 and AmyTracker680 (2 μg/mL). **g**, Representative image showing CHMP4B association about AmyTracker puncta after LLOMe exposure (1 mM, 10 minutes). **h**, (Left) schematic of experimental design and (right) representative images and quantitation of AmyTracker puncta presence in U-2 OS cells treated with LLOMe (1 mM, 10 minutes) followed by washout and further incubation for the indicated time (AmyTracker staining conducted as endpoint analysis); analysis of n ≥ 74 cells per replicate, standard deviation shown as error bars, comparisons analyzed using one-way ANOVA, ***=P≤0.001, **=P<0.01. Scalebars = 5 μm.

Here, we employ a breadth of *in vitro* and cell-based assays to conclude that (Leu-Leu)_n_ oligopeptides spontaneously assemble into membranolytic cross-β-sheet–rich amyloid fibrils within the lysosome. LLOMe analogs modulate the severity of lysosomal damage and cytotoxicity. We additionally report a second action of dipeptide esters: the induction of dipeptidyl ligase activity of CTSC onto proximal proteins, here termed Lysosome Injury-Localized Addition by Cathepsin C (LILAC).

## (Leu-Leu)_n_ assembles into amyloid fibrils

Previous characterization of (Leu-Leu)_n_ ligation products lacked structural analysis. Due to their high hydrophobicity and β-strand propensity, we hypothesized that (Leu-Leu)_n_ oligopeptides would spontaneously self-assemble into cross-β-sheet rich structures. Combining CTSC and LLOMe in a phosphate buffer (pH 6.5) resulted in the formation of a solid precipitate. Liquid chromatography–mass spectrometry (LC-MS) analysis of the DMSO-solubilized precipitate revealed the presence of (Leu-Leu)_1-4_ oligopeptides, primarily (Leu-Leu)_3_ (Fig. 1b), in general agreement with the previous study^29^. To probe structural order within the precipitate, we conducted an X-ray diffraction study, revealing diffraction rings at 11.5 and 4.8 Å (Fig. 1c), hallmarks of a cross-β-sheet amyloid fibril structure^32^. Additionally, negative stain electron microscopy revealed rod-like assemblies in the reaction mixture (Fig. 1d). Collectively, these results suggest that CTSC-catalyzed ligation of LLOMe affords (Leu-Leu)_3-4_ oligopeptides, that, upon exceeding some critical concentration and chain length, rapidly self-assemble into cross-β-sheet–rich amyloid fibrils (Fig. 1e).

In U-2 OS cells, LLOMe treatment resulted in visibly expanded LAMP1-positive endolysosomes (Extended Data Fig. 1a), possibly due to the genesis of lumenal amyloid fibrils. To probe for amyloid content within endolysosomes, cells were stained with the amyloid-binding dye AmyTracker^33^, resulting in robust AmyTracker fluorescence within endolysosomes (Fig. 1f). Alternative amyloid-binding fluorophores thioflavin T and Proteostat stained similarly (Extended Data Fig. 1b), consistent with amyloid fibrils within the lysosome. Lysosome deacidification by bafilomycin A1 was not sufficient to induce AmyTracker fluorescence (Extended Data Fig 1c), ruling out an elevated pH-dependent aggregation of endogenous lysosomal cargoes. Pretreatment with the CTSC inhibitor AZD5248, which forms a covalent bond with the active site residue Cys234^34^, ablated LLOMe-induced AmyTracker staining (Extended Data Fig. 1c), confirming CTSC activity is required to generate AmyTracker-positive species. AmyTracker puncta were consistently encircled by CHMP4B, a marker for ESCRT-mediated lysosomal repair^7,9^ (Fig. 1g, Extended Data Fig. 1d), suggesting that amyloid formation causes lysosomal damage. Overall, both *in vitro* and in-cell evidence support the spontaneous intralysosomal amyloid fibril formation of LLOMe-derived (Leu-Leu)_n_ oligopeptides. Previous demonstrations of amyloid fibril–membrane interactions^35-37^ and the observed near-universal association of (Leu-Leu)_n_ amyloid deposits with CHMP4B puncta together suggest (Leu-Leu)_n_ fibrils actively mediate lysosomal membranolysis.

### (Leu-Leu)_n_ fibril deposits are long lived

To model lysosomal repair, LLOMe-induced lysosomal damage has been carried out ‘reversibly’ by washing out LLOMe after a given treatment period^9,38^. Accordingly, we hypothesized that (Leu-Leu)_n_ amyloid fibrils would be rapidly degraded. In agreement with previous results^7^, we observed that after a 10-minute LLOMe treatment, CHMP4B was rapidly recruited to endolysosomes and dissipated within 3 hours after LLOMe washout (Extended Data Fig. 1e); however, endolysosomes remained visibly swollen at this timepoint. Strikingly, six hours after LLOMe washout, cells were still robustly AmyTracker positive, with puncta dimly present even after 12 hours (Fig. 1h). These results indicate the persistence of (Leu-Leu)_n_ amyloid fibrils, with fibril digestion occurring over tens of hours. The pathway(s) mediating fibril digestion were not explored in this study. Additionally, it remains unclear whether the observed fibril deposits occupy once-damaged then repaired lysosomes, or terminally damaged lysosomal vesicles destined for lysophagy^26^.

### CryoET observations of lysosomal fibrils

To provide additional evidence for intralysosomal amyloid fibril formation following LLOMe treatment, and to evaluate associated lysosomal membrane integrity, we conducted cryo-electron tomography studies employing a strategy to affinity isolate lysosomes (Extended Data Fig. 2).

HEK293T cells stably expressing the lysosomal membrane protein TMEM192-GFP, either untreated or treated with LLOMe for 10 minutes, were lysed by mechanical disruption that preserves lysosome integrity. Lysates were then incubated on anti-GFP nanobody-functionalized grids before plunge freezing and tilt series collection. Untreated lysosomes (from previously collected data provided by R. Woldeyes, H. Siems, B. Hill et al. available through the CryoET Data Portal^39^ Dataset ID: DS-10444) were intact and displayed no obvious lumenal material outside of intralumenal vesicles, consistent with other studies employing this lysosome enrichment methodology^40,41^ (Fig. 2, additional examples found in Extended Data Fig. 2). In stark contrast, a subset of lysosomes from LLOMe-treated cells bore fibrous cargo within either intact, partially damaged, or fully ruptured membranes. Overall, this observation provides further evidence that LLOMe treatment results in fibril formation within the lysosome associated with membranolysis.

**Fig. 2.**
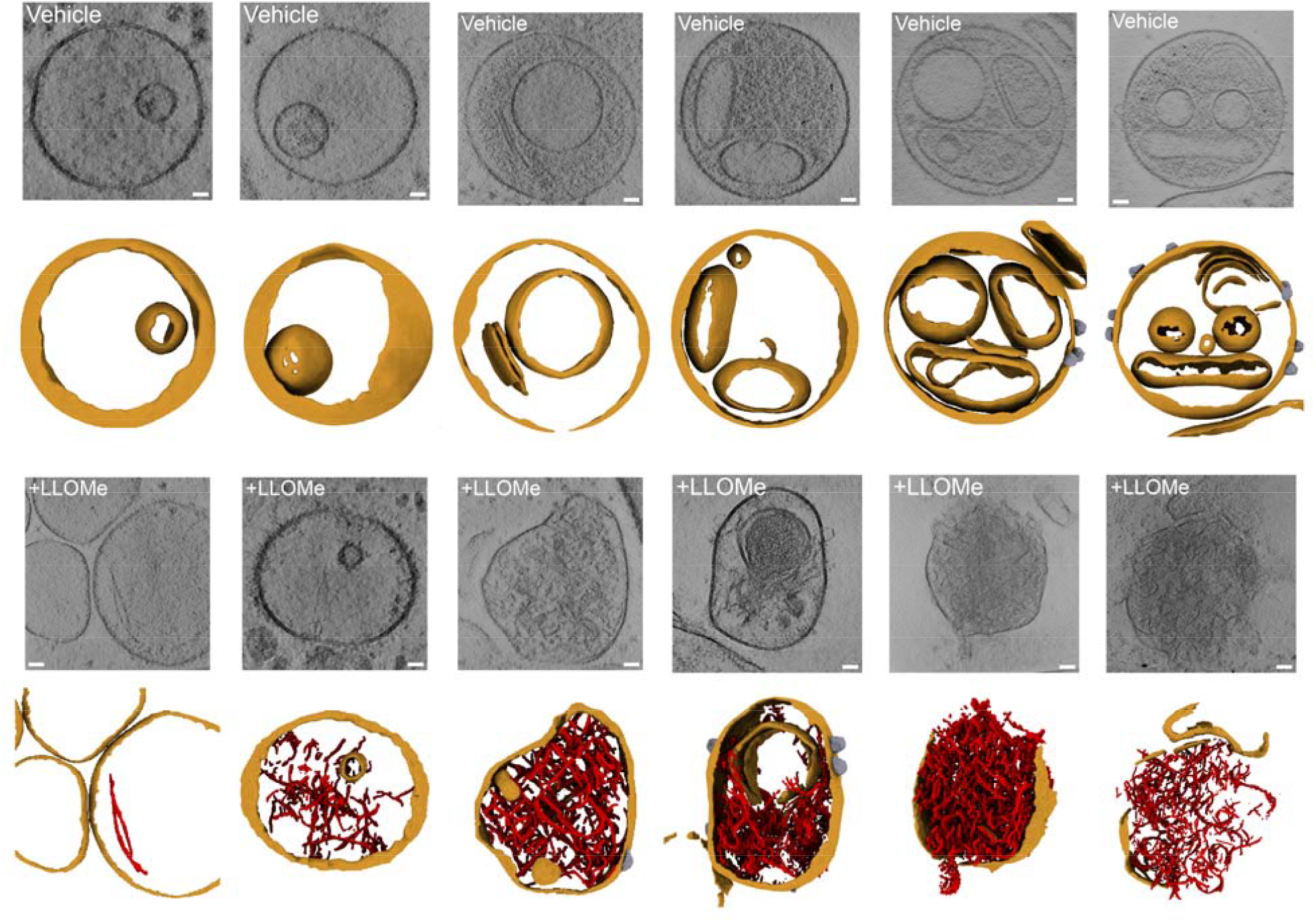
Cryo-electron tomography of affinity-captured lysosomes reveals membrane damage and fibril deposition. Representative tomographic slabs (average of five consecutive slices) and corresponding 3D segmentations of lysosomes from untreated HEK293T cells (top row) or LLOMe-treated cells (1 mM, 10 min; bottom row). In the segmentations, membranes are colored orange, fibrils red, and flotillin structures gray. Scale bars =10 nm

### Dipeptide ester sequence variation

We anticipated that dipeptide ester sequence variation could alter cellular phenotypes by altering resultant CTSC-generated oligopeptide assembly structure and membrane interactions.

Phenylalanine was utilized as an aromatic, hydrophobic residue that is independently amyloidogenic^42^ in three dipeptide esters: LF-, FL-, and FFOMe (Fig. 3a). Additionally, we synthesized a propargyl dipeptide termed ULOMe (Fig. 3a) as a representative dipeptide containing a noncanonical amino acid with potential for downstream bioorthogonal chemistry (molecular docking studies suggested an optimal carbon chain length of the propargyl sidechain that preserved productive engagement with CTSC (Extended Data Fig. 3a)). Tetrapeptide ligation products were observed for all dipeptides upon incubation with CTSC *in vitro* (Extended Data Fig. 3b-e). In U-2 OS cells, we observed LC3B-II accumulation, a marker of lysophagy^26^ and ATG8-mediated lysosomal damage response^27,43^, with all dipeptides studied (Fig. 3b).

**Fig. 3.**
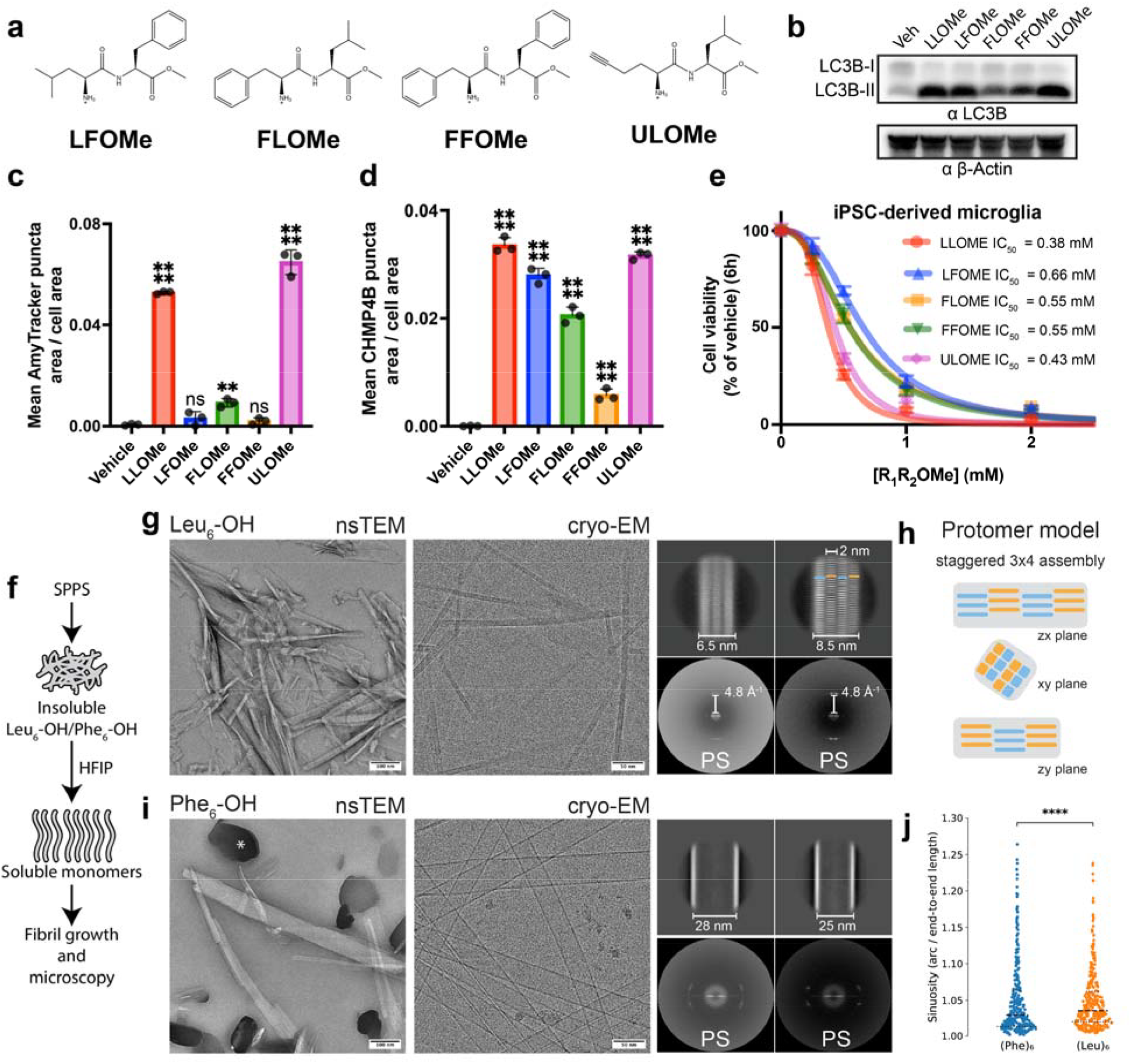
LLOMe analog modulation of cellular phenotypes is linked with amyloidogenicity of oligopeptide fibril structures. **a**, Structures of LLOMe analogs utilized here. **b**, Western blot displaying accumulation of LC3B-II upon treatment of the indicated dipeptide methyl esters (1 mM, 60 minutes). Quantitation of **c**, AmyTracker or **d**, CHMP4B puncta induced upon treatment with the indicated dipeptide esters; analysis of n ≥ 55 cells per replicate per timepoint, standard deviation shown as error bars, comparisons against vehicle treatment analyzed using one-way ANOVA, ****=P≤0.0001, **=P<0.01, ns=P>0.05. **e**, Viability plot of iPSC-derived microglial cultures against 6-hour dipeptide ester exposure of the indicated concentration; each point represents the mean of triplicate measurements, standard deviation shown as error bars where possible, data fit to variable slope IC_50_ curves. **F**, Schematic of Leu_6_ and Phe_6_ preparation. **g**, negative-stain and cryo-EM micrographs of Leu_6_, 2D class averages and power spectra (PS) with layer-line annotation at 4.8 Å^-1^. **h**, Staggered 3×4 protomer model of Leu_6_ along the zx, xy and zy planes. **i**, negative-stain and cryo-EM micrographs of Phe_6_, 2D class averages and power spectra; asterisk indicates uranyl formate crystal. **d**, Fibril sinuosity comparison (ratio of arc over end-to-end fibril length). Mann-Whitney U gives z = −29.8, p = 1.5×10 □^1^ □ □, nPhe_6_ = 372,441 and nLeu_6_ = 122,018.

Endolysosomes were visibly enlarged for all treatments; however, LF-, FL-, and FFOMe were not detectably stained by AmyTracker (Fig. 3c, Extended Data Fig. 4a), but were instead readily stained by Proteostat (Extended Data Fig. 4b) suggesting divergence in oligopeptide assembly structure relative to LLOMe. In contrast, ULOMe exhibited dye fluorescence analogous to LLOMe (Fig. 3c). Total CHMP4B puncta formation, a proxy for totality of lysosomal permeabilization, was similar between LLOMe and ULOMe (Fig. 3d, Extended Data Fig. 4c), but diminished with LF-, FL-, and FFOMe, with FFOMe inducing the weakest CHMP4B response. Analysis in HEK293T confirmed that AmyTracker binding or lack thereof was dipeptide sequence dependent and further supported FFOMe as a relatively weak inducer of the ESCRT repair response (Extended Data Fig. 5).

Lysosomal damage-mediated cell death^44^ was characterized utilizing LL-, LF-, FL-, FF-, and ULOMe in iPSC-derived microglia, which exhibit high lysosomal function^45,46^ and are susceptible to lysosomal damage-mediated cell death^29^. A major loss in viability was observed after only a 6-hour exposure of 0.5 mM LLOMe, resulting in an IC_50_ of 0.38 mM (Fig. 3e).

LFOMe, FLOMe, and FFOMe displayed reduced cytotoxicities, with IC_50_’s of 0.66, 0.55, and 0.55 mM, respectively. ULOMe again behaved similarly to LLOMe with an IC50 of 0.43 mM, displaying an association between toxicity and the degree of induced lysosomal membrane damage.

We hypothesized that the degree of lysosome membranolysis was dependent on oligopeptide assembly structure. To investigate this, we utilized solid-phase peptide synthesis to produce multimilligram quantities of the hexapeptides Leu_6_-OH and Phe_6_-OH, utilizing the carboxylate to enhance solubility. After synthesis, potential aggregates were monomerized using Hexafluoroisopropanol and dried over a stream of N_2_ gas before resolubilization in DMSO. Fibrils were grown by diluting 2 µM hexapeptide in HEPES buffer (pH 6.5) for four days before imaging using negative stain TEM and cryo-EM (Fig. 3f). Leu_6_ fibrils displayed low to no twist, preventing successful helical reconstruction, but the fast Fourier transform (power spectrum) of well-aligning 2D class averages showed layer lines at the 4.8 Å^-1^ frequency, indicative of a cross-β-sheet structure (Fig. 3g). Additionally, different 2D averages show staggered protomers, potentially assembling into 3×4 fibrils with protomers about 2 nm in length (Fig. 3g, h). Phe_6_ formed hollow tubular assemblies with a wide range of tube diameters, visible on a per-micrograph level and in 2D class averages (Fig. 3i), where well-aligning segments assembled into tubes of either 25 nm or 28 nm diameters. For these tubes, a clear amyloid-specific layer line is absent in the power spectrum, but off-meridian reflections are observable, indicative of a clear helical assembly. Leu_6_ fibrils show significantly higher sinuosity compared to Phe_6_ fibrils (Fig. 3j), indicating higher fibril bending for the thinner Leu_6_ compared to the wider nanotubes of Phe_6_. Collectively, the data suggests that efficient lysosome membranolysis is carried out by (Leu-Leu)_n_ through assembly into cross-β-sheet rich protofibril building blocks that we hypothesize further laterally assemble to carpet the membrane surface, inducing membrane tears through active assembly (Fig. 3g)^25,47^. Notably, Phe_6_ assemblies lack cross-β-sheet structure and protofibril building blocks, which may account for the reduced lysosome-damaging phenotypes observed employing FFOMe.

### Inducible proximity labeling by CTSC

While CTSC ligase activity has been characterized as a minor reaction at elevated pH *in vitro* in this and other studies^12,13^, intralysosomal ligation of LLOMe into oligopeptides suggests ligase activity may be more prevalent in biological contexts than previously appreciated. Under saturating conditions, CTSC predominantly exists as the ligation-competent dipeptidyl thioester intermediate^48^. We observed that CTSC ligation not only generates homo-oligomers from a given starting material but can generate heterologous products from a peptide mixture (Extended Data Fig. 6a-c). Low selectivity for incoming peptide nucleophiles is supported by the absence of well-defined S1’ and S2’ pockets in the CTSC structure^49^. We posited that the dipeptidyl-CTSC thioester intermediate formed by the presence of excess dipeptide ester could be subject to nucleophilic attack by a wide-range of proximal peptide and protein nucleophiles, resulting in the formation of distinct dipeptidyl-adducts (Extended Data Fig. 6d). To investigate this we employed ULOMe, a dipeptide containing a terminal alkyne amenable to Copper-catalyzed Azide–Alkyne Cycloaddition (CuAAC) reactions^50^, where addition of azide-bearing reporter molecules permits direct detection of UL-ligated species.

Cells were treated with increasing concentrations of ULOMe, lysed, and Rhodamine-N_3_ was added by CuAAC to visualize UL-adducts by SDS-PAGE in-gel fluorescence. Fluorescence intensity increased with ULOMe concentration throughout the entire lane and in multiple specific bands ranging from ∼5-150 kDa (Fig. 4a). The majority of fluorescence intensity was eliminated upon pretreatment with CTSC inhibitor AZD5248 or pan-cysteine cathepsin inhibitor E-64d, indicating that the appearance of these bands requires CTSC activity. Additionally, pretreatment with bafilomycin A1 did not increase or decrease overall signal intensity, suggesting UL-dipeptidyl ligation may be independent of initial lysosomal pH. To further support the necessity of active CTSC, we utilized a HeLa CTSC knockout line, which displayed neither punctate AmyTracker staining nor CHMP4B puncta following ULOMe treatment (Fig. 4b). UL-dipeptidyl ligation was eliminated in knockout cells (Fig. 4c). Overexpression in knockout cells of wildtype CTSC, but not a catalytically inactive C234A mutant, rescued UL-dipeptidyl ligation. Comparison of UL-adduct formation in a small panel of cell lines differentially expressing CTSC revealed relative signal is not consistently proportional to CTSC expression (Fig. 4d), suggesting accessible CTSC ligase activity, not solely CTSC expression, is responsible for UL-dipeptidyl ligation. Furthermore, we observed that the majority of UL-ligation occurs within 60 minutes, where prolonged incubations did not proportionately increase fluorescent signal likely owing to non-CTSC, cytosolic esterase activity depletion of ULOMe.

**Fig. 4.**
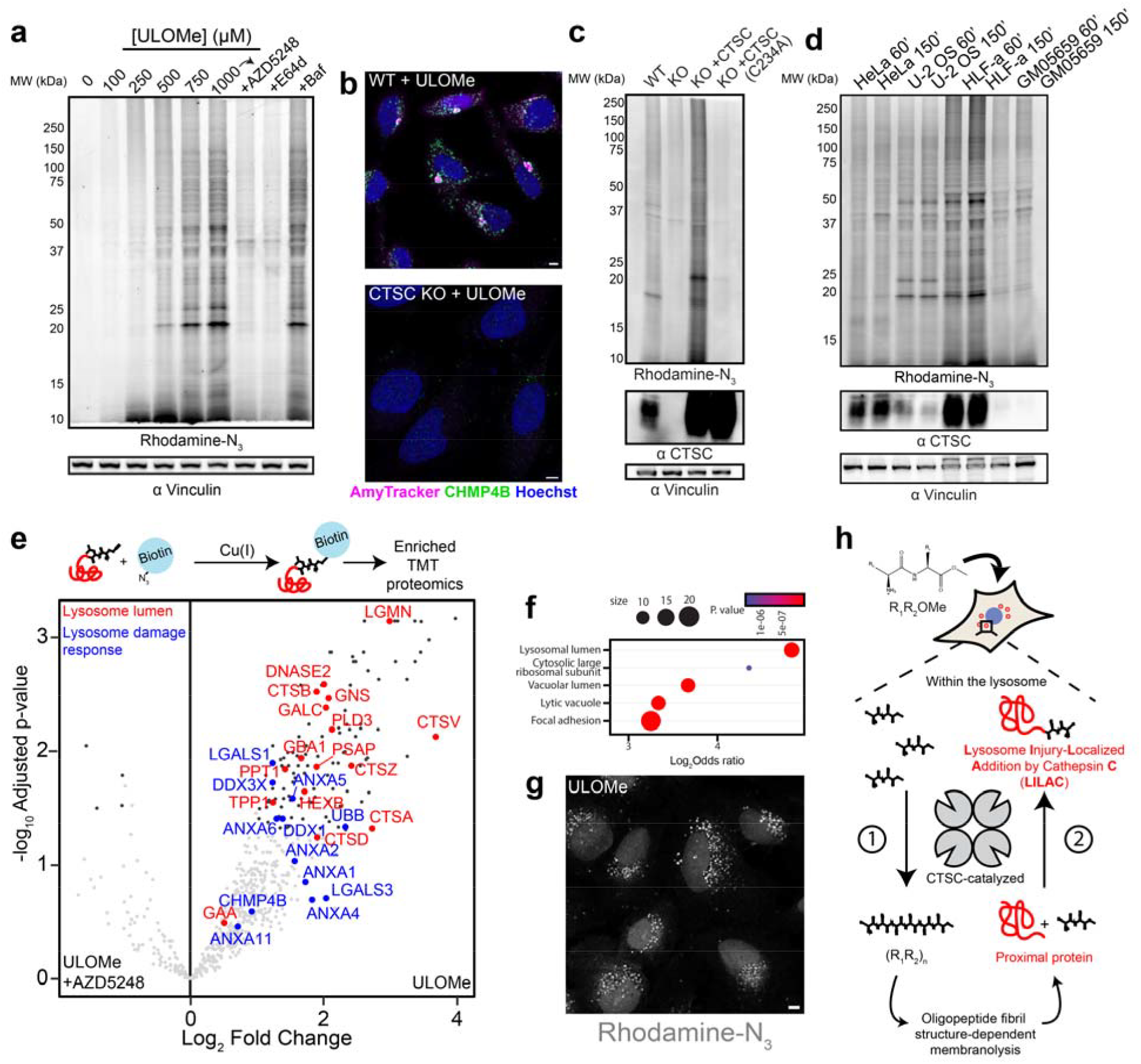
Dipeptide esters induce nonspecific dipeptidyl ligase activity by CTSC on distinct, proximal proteins. **a**, SDS-PAGE in-gel fluorescence of U-2 OS cell lysate after treatment with indicated concentrations of ULOMe (60 minutes) or 1 mM ULOMe after 90-minute pretreatment with CTSC inhibitor AZD5248 (10 μM) or pan-cysteine cathepsin inhibitor E-64d (10 μM), followed by addition of rhodamine-N_3_ by CuAAC in lysates. **b**, Representative images displaying AmyTracker and CHMP4B puncta formation induced by ULOMe (1 mM, 60 minutes) in WT, but not CTSC KO HeLa cells. **c**, In-gel fluorescence and CTSC and vinculin western blots of WT, KO, and KO HeLa cells overexpressing WT CTSC or CTSC(C234A) treated with ULOMe (1 mM, 60 minutes). **d**, In-gel fluorescence and CTSC and vinculin western blots of HeLa, U-2 OS, HLF-a, and GM05659 cells treated with ULOMe (1 mM, 60 or 150 minutes). **e**, (Top) brief schematic of proteomics sample preparation, (bottom) volcano plot for ULOMe treatment (1 mM, 60 minutes) with active versus inhibited CTSC, annotated with known lysosome lumen (red) and damage response (blue) proteins. **f**, GO enrichment (cellular compartment) for proteins enriched with active CTSC. **g**, Representative image visualizing UL-species localization after addition of Rhodamine-N_3_ by CuAAC. **h**, Model of dipeptide ester activity in cells: certain dipeptide esters are polymerized by CTSC into oligopeptides with propensity to form fibrillar strictures. Lysosome membrane permeabilization ensues dependent on fibril structural properties, activating nonspecific ligase activity of CTSC on proximal proteins, an activity termed Lysosome Injury-Localized Addition by Cathepsin C (LILAC). Scalebars = 5 μm

Investigating the effects of amino acid substitutions, we found that UFOMe, which behaved similar to LFOMe (Extended data Fig. 7a-c), also formed adducts with a broad range of proteins, although to a relatively lower extent than ULOMe (Extended Data Fig. 7d). We hypothesized the extent of dipeptide ester-induced membrane damage and adduct formation are coupled. To scrutinize this, we employed UYOMe, a non-damaging dipeptide ester (Extended Data Fig.7e, f). Employing UYOMe alone, UY-adducts were not observable. However, upon 10-minute preincubation with LLOMe following addition of UYOMe, UY-adducts became visible (Extended Data Fig.7g). In contrast, treatments using lysosome deacidification agents bafilomycin A1, chloroquine, or nigericin with UYOMe did not induce UY-adduct formation, suggesting that membrane damage, not just lysosome deacidification, may provide a signal which activates biological CTSC ligase activity, although this merits further mechanistic interrogation.

To investigate UL-adduct identities, we employed quantitative biotin-enriched proteomics. After treatment of cells in biological quadruplicate with DMSO, in triplicate with ULOMe alone, or in triplicate with ULOMe after preincubation of CTSC inhibitor AZD5248, biotin-N_3_ was added to UL-adducts by CuAAC, and samples were processed for biotin enrichment and 10-plex Tandem Mass Tagging (TMT)-based proteomics (Fig. 4e, Supplementary Table 1). Comparing ULOMe treatment without and with CTSC inhibition, of the 650 proteins identified, 128 were significantly enriched with active CTSC (Log_2_FC > 1; p. adj < 0.05). Several lysosomal hydrolases including GBA1, DNASE2, and PLD3 were identified, driving enrichment of the lysosome lumen in GO cellular component analysis (Fig. 4f). Additionally enriched were lysosome damage response proteins, including markers of lysophagy (LGALS1, UBB), stress granules (DDX3X, DDX1), and annexin family recruitment^11,51,52^. We speculate additional identified proteins are either undigested autophagy substrates or take part in as-of-yet uncharacterized facets of lysosome biology. Imaging revealed accumulation of UL-species in discrete perinuclear puncta dependent on CTSC activity, supporting the spatial specificity of UL-ligation (Fig. 4g, Extended Data Fig.8).

Altogether, we propose the following mechanistic model of dipeptide ester function: initially and within the lysosome, certain dipeptide esters are ligated by CTSC into oligopeptides predisposed to self-assemble into fibrillar structures, where cross-β-sheet amyloid fibril assemblies are exceptionally efficient at membrane disruption. Secondarily and apparently only in sufficiently damaged lysosomes, CTSC then catalyzes the ligation of excess dipeptide onto proximal proteins, an activity we term Lysosome Injury-Localized Addition by Cathepsin C (LILAC) (Fig. 4h).

## Discussion

To our knowledge, LLOMe represents the only reported reagent that rapidly induces the intralysosomal generation of amyloid fibrils, as reported herein. This observation was concurrently verified by a separate, independent investigation conducted in parallel^53^. While historic and current use of LLOMe focuses on the rapid lysosomal damage phenotype, here we observed evidence of persistent deposits of LLOMe-derived fibrils tens of hours after treatment (Fig. 1h). In light of this, we propose LLOMe and a subset of analogs represent a generalizable model system for studying mechanisms governing intracellular amyloid fibril formation, membrane interactions, and degradation kinetics in biological systems relating to neurodegeneration.

Proximity labeling has established itself as a powerful approach to profile the local environment of given protein of interest^54-56^. Here we demonstrate an unexpected consequence of dipeptide ester treatment: CTSC catalyzes the ligation of certain dipeptides onto proximal proteins, effectively acting as a proximity labeling enzyme in an activity we term LILAC. In cells, LILAC appears restricted to damaged lysosomes, yet once activated, is generally agnostic to the identity of the incoming protein nucleophile implying a potential unexplored role of CTSC in lysosome biology. Exploiting LILAC facilitated the proximity-based proteomics study of the damaged lysosome lumen, a notably inaccessible locale for conventional proximity-labeling enzyme fusion approaches. To our knowledge, use of a fully endogenous, non-engineered enzyme for spatially localized proximity-labeling has not been previously reported. We envision this lowered experimental barrier, contrasted to the relatively heavier genetic engineering or chemical synthesis load required for other proximity-labeling approaches, to facilitate proximity-based proteomics and imaging studies of the damaged lysosome in any feasible biological system endogenously expressing CTSC. We anticipate this work to spur the development of novel chemical substrates exploiting other endogenous enzymes for proximity labeling within discrete organelles and in various other relevant locales.

## Data and materials availability

The Cryo-electron tomography dataset derived from LLOMe-treated cells collected here is available on the CZ CryoET Data Portal (*deposition_number TBA*). The presented control dataset was provided by R. Woldeyes, H. Siems, B. Hill et al. and is available through CryoET Data Portal^39^ Dataset ID: DS-10444. Relevant data from image analysis is available upon request. LFOMe, FLOMe, ULOMe, UFOMe, and UYOMe materials are available from the Kelly lab under the terms of an MTA.

## Acknowledgments

We acknowledge the expert assistance of Scott Henderson, Kimberly Vanderpool, and Theresa Fassel of The Core Microscopy Facility at The Scripps Research Institute. We acknowledge the expert assistance of Jake Bailey at the UCSD X-ray Crystallography Facility. We thank Emily P. Bentley for providing expert editorial assistance. Research reported in this publication was supported by the National Institute on Aging of the National Institutes of Health grant RF1AG073418 (JWK) and the National Center for Advancing Translational Sciences of the National Institutes of Health grant T32TR004396 (RDE). The content is solely the responsibility of the authors and does not necessarily represent the official views of the National Institutes of Health. Funding for this research was provided by the German Research Council project number 556478029 to JHS. RTO is thankful for the financial support of the George E. Hewitt Foundation for Medical Research. Further support was provided by the Department of Education under the Title V DHSI Program Grant #P031S240270 (DAY). Additional funding was provided by the Freedom Together Foundation (JWK). The CZ Imaging Institute is fully funded by the Chan Zuckerberg Initiative (CZII-2023–327779).

## Author contributions

Conceptualization: RDE and JWK

Methodology: RDE, SA, and DS

Investigation: RDE, SA, IID, RTO, JHS, HS, EAM, UHE, CA, FA, DAY, OLL, DS

Funding acquisition: BFC, DAA, JWK

Supervision: DAA, BFC, JWK

Writing – original draft: RDE

Writing – review & editing: RDE, JWK

## References

1 Pryor, P. R. & Luzio, J. P. Delivery of endocytosed membrane proteins to the lysosome. Biochimica et Biophysica Acta (BBA) - Molecular Cell Research 1793, 615–624 (2009). 10.1016/j.bbamcr.2008.12.022

2 Nixon, R. A. & Rubinsztein, D. C. Mechanisms of autophagy–lysosome dysfunction in neurodegenerative diseases. Nature Reviews Molecular Cell Biology 25, 926–946 (2024). 10.1038/s41580-024-00757-5

3 Rose, K. et al. Tau fibrils induce nanoscale membrane damage and nucleate cytosolic tau at lysosomes. Proceedings of the National Academy of Sciences 121, e2315690121 (2024). 10.1073/pnas.2315690121

4 Lee, J.-H. et al. Faulty autolysosome acidification in Alzheimer’s disease mouse models induces autophagic build-up of Aβ in neurons, yielding senile plaques. Nature Neuroscience 25, 688–701 (2022). 10.1038/s41593-022-01084-8

5 Chou, C.-C. et al. Proteostasis and lysosomal repair deficits in transdifferentiated neurons of Alzheimer’s disease. Nature Cell Biology 27, 619–632 (2025). 10.1038/s41556-025-01623-y

6 Chen, J. J. et al. Compromised function of the ESCRT pathway promotes endolysosomal escape of tau seeds and propagation of tau aggregation. Journal of Biological Chemistry 294, 18952–18966 (2019). 10.1074/jbc.RA119.009432

7 Radulovic, M. et al. ESCRT-mediated lysosome repair precedes lysophagy and promotes cell survival. The EMBO Journal 37, e99753 (2018). 10.15252/embj.201899753

8 Radulovic, M. et al. Cholesterol transfer via endoplasmic reticulum contacts mediates lysosome damage repair. The EMBO Journal 41, e112677 (2022). 10.15252/embj.2022112677

9 Skowyra, M. L., Schlesinger, P. H., Naismith, T. V. & Hanson, P. I. Triggered recruitment of ESCRT machinery promotes endolysosomal repair. Science 360, eaar5078 (2018). doi:10.1126/science.aar5078

10 Tan, J. X. & Finkel, T. A phosphoinositide signalling pathway mediates rapid lysosomal repair. Nature 609, 815–821 (2022). 10.1038/s41586-022-05164-4

11 Eapen, V. V., Swarup, S., Hoyer, M. J., Paulo, J. A. & Harper, J. W. Quantitative proteomics reveals the selectivity of ubiquitin-binding autophagy receptors in the turnover of damaged lysosomes by lysophagy. eLife 10, e72328 (2021). 10.7554/eLife.72328

12 Heinrich, C. P. & Fruton, J. S. The action of dipeptidyl transferase as a polymerase. Biochemistry 7, 3556–3565 (1968). 10.1021/bi00850a033

13 Würz, H., Tanaka, A. & Fruton, J. S. Polymerization of Dipeptide Amides by Cathepsin C*. Biochemistry 1, 19–29 (1962). 10.1021/bi00907a004

14 Root, J., Merino, P., Nuckols, A., Johnson, M. & Kukar, T. Lysosome dysfunction as a cause of neurodegenerative diseases: Lessons from frontotemporal dementia and amyotrophic lateral sclerosis. Neurobiology of Disease 154, 105360 (2021). 10.1016/j.nbd.2021.105360

15 Nixon, R. A. Amyloid precursor protein and endosomal-lysosomal dysfunction in Alzheimer’s disease: inseparable partners in a multifactorial disease. Faseb j 31, 2729–2743 (2017). 10.1096/fj.201700359

16 Wallings, R. L., Humble, S. W., Ward, M. E. & Wade-Martins, R. Lysosomal Dysfunction at the Centre of Parkinson’s Disease and Frontotemporal Dementia/Amyotrophic Lateral Sclerosis. Trends in Neurosciences 42, 899–912 (2019). 10.1016/j.tins.2019.10.002

17 Zhang, W. et al. Impairment of the autophagy–lysosomal pathway in Alzheimer’s diseases: Pathogenic mechanisms and therapeutic potential. Acta Pharmaceutica Sinica B 12, 1019–1040 (2022). 10.1016/j.apsb.2022.01.008

18 Udayar, V., Chen, Y., Sidransky, E. & Jagasia, R. Lysosomal dysfunction in neurodegeneration: emerging concepts and methods. Trends in Neurosciences 45, 184–199 (2022). 10.1016/j.tins.2021.12.004

19 Ayers, J. I., Giasson, B. I. & Borchelt, D. R. Prion-like Spreading in Tauopathies. Biol Psychiatry 83, 337–346 (2018). 10.1016/j.biopsych.2017.04.003

20 Jan, A., Gonçalves, N. P., Vaegter, C. B., Jensen, P. H. & Ferreira, N. The Prion-Like Spreading of Alpha-Synuclein in Parkinson’s Disease: Update on Models and Hypotheses. Int J Mol Sci 22 (2021). 10.3390/ijms22158338

21 Flavin, W. P. et al. Endocytic vesicle rupture is a conserved mechanism of cellular invasion by amyloid proteins. Acta Neuropathologica 134, 629–653 (2017). 10.1007/s00401-017-1722-x

22 Kakuda, K. et al. Lysophagy protects against propagation of α-synuclein aggregation through ruptured lysosomal vesicles. Proceedings of the National Academy of Sciences 121, e2312306120 (2024). doi:10.1073/pnas.2312306120

23 Yuste-Checa, P. et al. The extracellular chaperone Clusterin enhances Tau aggregate seeding in a cellular model. Nature Communications 12, 4863 (2021). 10.1038/s41467-021-25060-1

24 Kolay, S. et al. The dual fates of exogenous tau seeds: Lysosomal clearance versus cytoplasmic amplification. Journal of Biological Chemistry 298 (2022). 10.1016/j.jbc.2022.102014

25 Sciacca, M. F. M., La Rosa, C. & Milardi, D. Amyloid-Mediated Mechanisms of Membrane Disruption. Biophysica 1, 137–156 (2021).

26 Maejima, I. et al. Autophagy sequesters damaged lysosomes to control lysosomal biogenesis and kidney injury. The EMBO Journal 32, 2336–2347 (2013). 10.1038/emboj.2013.171

27 Cross, J. et al. Lysosome damage triggers direct ATG8 conjugation and ATG2 engagement via non-canonical autophagy. Journal of Cell Biology 222 (2023). 10.1083/jcb.202303078

28 Jia, J. et al. Stress granules and mTOR are regulated by membrane atg8ylation during lysosomal damage. Journal of Cell Biology 221 (2022). 10.1083/jcb.202207091

29 Thiele, D. L. & Lipsky, P. E. Mechanism of L-leucyl-L-leucine methyl ester-mediated killing of cytotoxic lymphocytes: dependence on a lysosomal thiol protease, dipeptidyl peptidase I, that is enriched in these cells. Proc Natl Acad Sci U S A 87, 83–87 (1990). 10.1073/pnas.87.1.83

30 McGuire, M. J., Lipsky, P. E. & Thiele, D. L. Purification and characterization of dipeptidyl peptidase I from human spleen. Arch Biochem Biophys 295, 280–288 (1992). 10.1016/0003-9861(92)90519-3

31 Uchimoto, T. et al. Mechanism of apoptosis induced by a lysosomotropic agent, L-Leucyl-L-Leucine methyl ester. Apoptosis 4, 357–362 (1999). 10.1023/a:1009695221038

32 Li, H. et al. in Encyclopedia of Analytical Chemistry (2009).

33 Pretorius, E. et al. Both lipopolysaccharide and lipoteichoic acids potently induce anomalous fibrin amyloid formation: assessment with novel Amytracker™ stains. J R Soc Interface 15 (2018). 10.1098/rsif.2017.0941

34 Furber, M. et al. Cathepsin C Inhibitors: Property Optimization and Identification of a Clinical Candidate. Journal of Medicinal Chemistry 57, 2357–2367 (2014). 10.1021/jm401705g

35 Arispe, N., Rojas, E. & Pollard, H. B. Alzheimer disease amyloid beta protein forms calcium channels in bilayer membranes: blockade by tromethamine and aluminum. Proceedings of the National Academy of Sciences 90, 567–571 (1993). doi:10.1073/pnas.90.2.567

36 Lashuel, H. A., Hartley, D., Petre, B. M., Walz, T. & Lansbury, P. T. Amyloid pores from pathogenic mutations. Nature 418, 291–291 (2002). 10.1038/418291a

37 Viles, J. H. Imaging Amyloid-β Membrane Interactions: Ion-Channel Pores and Lipid-Bilayer Permeability in Alzheimer’s Disease. Angewandte Chemie International Edition 62, e202215785 (2023). 10.1002/anie.202215785

38 Gahlot, P. et al. Lysosomal damage sensing and lysophagy initiation by SPG20-ITCH. Molecular Cell 84, 1556–1569.e1510 (2024). 10.1016/j.molcel.2024.02.029

39 Ermel, U. et al. A data portal for providing standardized annotations for cryo-electron tomography. Nature Methods 21, 2200–2202 (2024). 10.1038/s41592-024-02477-2

40 Peck, A. et al. A realistic phantom dataset for benchmarking cryo-ET data annotation. Nature Methods 22, 1819–1823 (2025). 10.1038/s41592-025-02800-5

41 Peck, A. et al. AreTomoLive: automated reconstruction of comprehensively corrected and denoised cryo-electron tomograms in real time and at high throughput. Nature Methods (2026). 10.1038/s41592-026-03093-y

42 Adler-Abramovich, L. et al. Phenylalanine assembly into toxic fibrils suggests amyloid etiology in phenylketonuria. Nature Chemical Biology 8, 701–706 (2012). 10.1038/nchembio.1002

43 Nakamura, S. et al. LC3 lipidation is essential for TFEB activation during the lysosomal damage response to kidney injury. Nature Cell Biology 22, 1252–1263 (2020). 10.1038/s41556-020-00583-9

44 Serrano-Puebla, A. & Boya, P. Lysosomal membrane permeabilization in cell death: new evidence and implications for health and disease. Ann N Y Acad Sci 1371, 30–44 (2016). 10.1111/nyas.12966

45 Gao, C., Jiang, J., Tan, Y. & Chen, S. Microglia in neurodegenerative diseases: mechanism and potential therapeutic targets. Signal Transduction and Targeted Therapy 8, 359 (2023). 10.1038/s41392-023-01588-0

46 Majumdar, A. et al. Activation of Microglia Acidifies Lysosomes and Leads to Degradation of Alzheimer Amyloid Fibrils. Molecular Biology of the Cell 18, 1490–1496 (2007). 10.1091/mbc.e06-10-0975

47 Tian, Y., Liang, R., Kumar, A., Szwedziak, P. & Viles, J. H. 3D-visualization of amyloid-β oligomer interactions with lipid membranes by cryo-electron tomography. Chemical Science 12, 6896–6907 (2021). 10.1039/D0SC06426B

48 Schneck, J. L. et al. Chemical Mechanism of a Cysteine Protease, Cathepsin C, As Revealed by Integration of both Steady-State and Pre-Steady-State Solvent Kinetic Isotope Effects. Biochemistry 47, 8697–8710 (2008). 10.1021/bi8007627

49 Turk, D. et al. Structure of human dipeptidyl peptidase I (cathepsin C): exclusion domain added to an endopeptidase framework creates the machine for activation of granular serine proteases. The EMBO Journal 20, 6570–6582 (2001). 10.1093/emboj/20.23.6570

50 Hein, J. E. & Fokin, V. V. Copper-catalyzed azide-alkyne cycloaddition (CuAAC) and beyond: new reactivity of copper(I) acetylides. Chem Soc Rev 39, 1302–1315 (2010). 10.1039/b904091a

51 Bussi, C. et al. Stress granules plug and stabilize damaged endolysosomal membranes. Nature 623, 1062–1069 (2023). 10.1038/s41586-023-06726-w

52 Yim, W. W., Yamamoto, H. & Mizushima, N. Annexins A1 and A2 are recruited to larger lysosomal injuries independently of ESCRTs to promote repair. FEBS Lett 596, 991–1003 (2022). 10.1002/1873-3468.14329

53 Li, D. et al. Cathepsin-dependent amyloid formation drives mechanical rupture of lysosomal membranes. bioRxiv, 2026.2001.2017.700056 (2026). 10.64898/2026.01.17.700056

54 Rhee, H.-W. et al. Proteomic Mapping of Mitochondria in Living Cells via Spatially Restricted Enzymatic Tagging. Science 339, 1328–1331 (2013). doi:10.1126/science.1230593

55 Branon, T. C. et al. Efficient proximity labeling in living cells and organisms with TurboID. Nature Biotechnology 36, 880–887 (2018). 10.1038/nbt.4201

56 Roux, K. J., Kim, D. I., Raida, M. & Burke, B. A promiscuous biotin ligase fusion protein identifies proximal and interacting proteins in mammalian cells. J Cell Biol 196, 801–810 (2012). 10.1083/jcb.201112098

