## Supplementary Figures for "Intralysosomal Amyloidogenesis and Proximity Labeling by Cathepsin C"

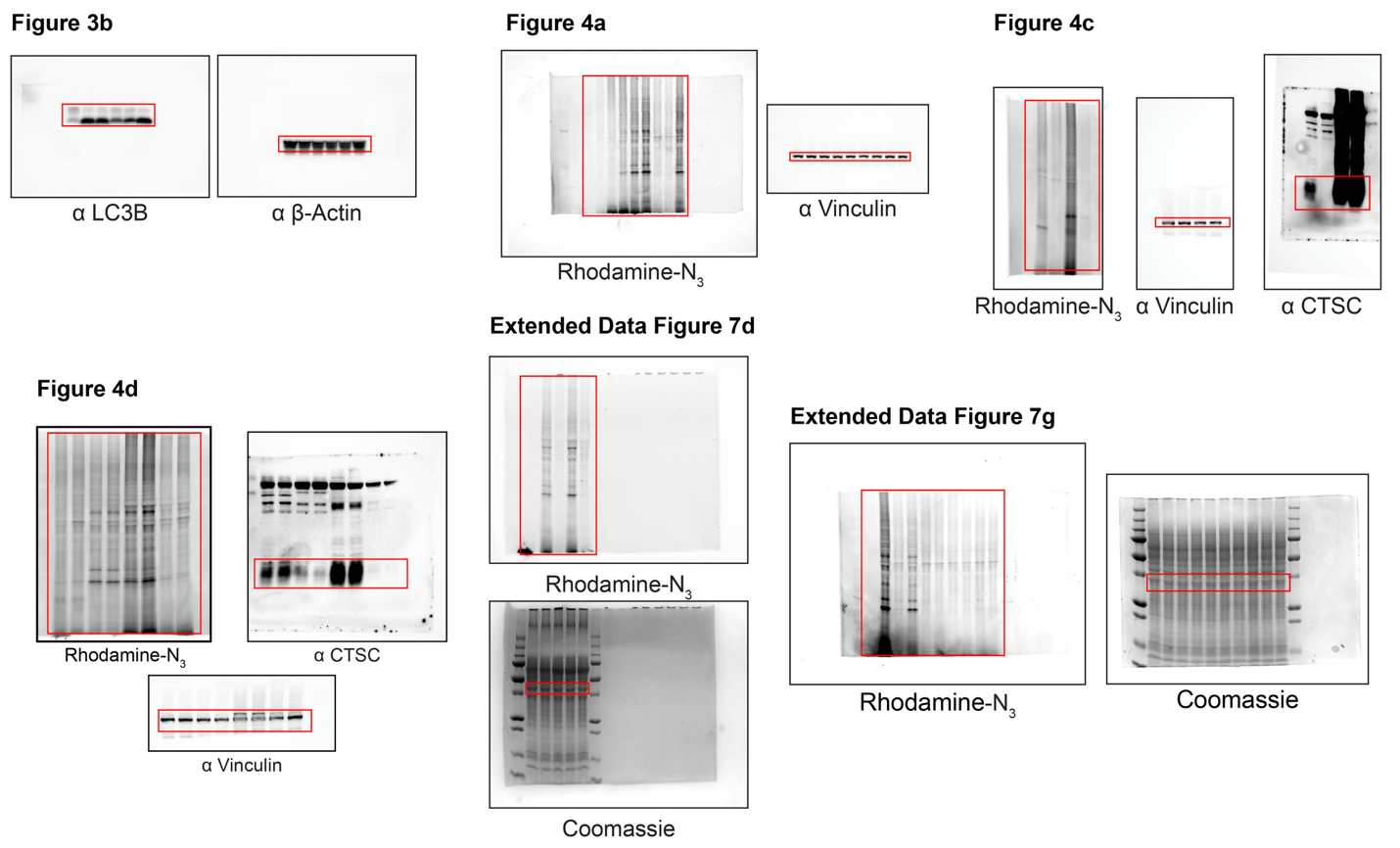

Supplementary Figure 1. Uncropped western blot and in-gel fluorescence images.

Parent figures are denoted, regions highlighted in red are presented in figures.

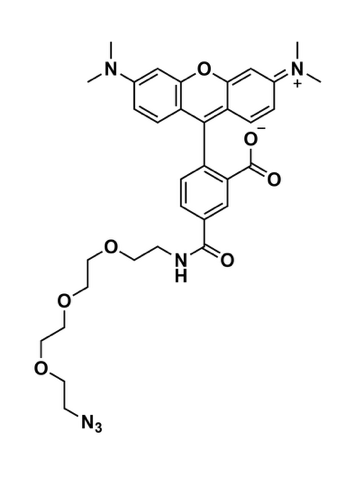

**Supplementary Figure 2. Structure of Rhodamine-PEG_3_-Azide used in this study.**

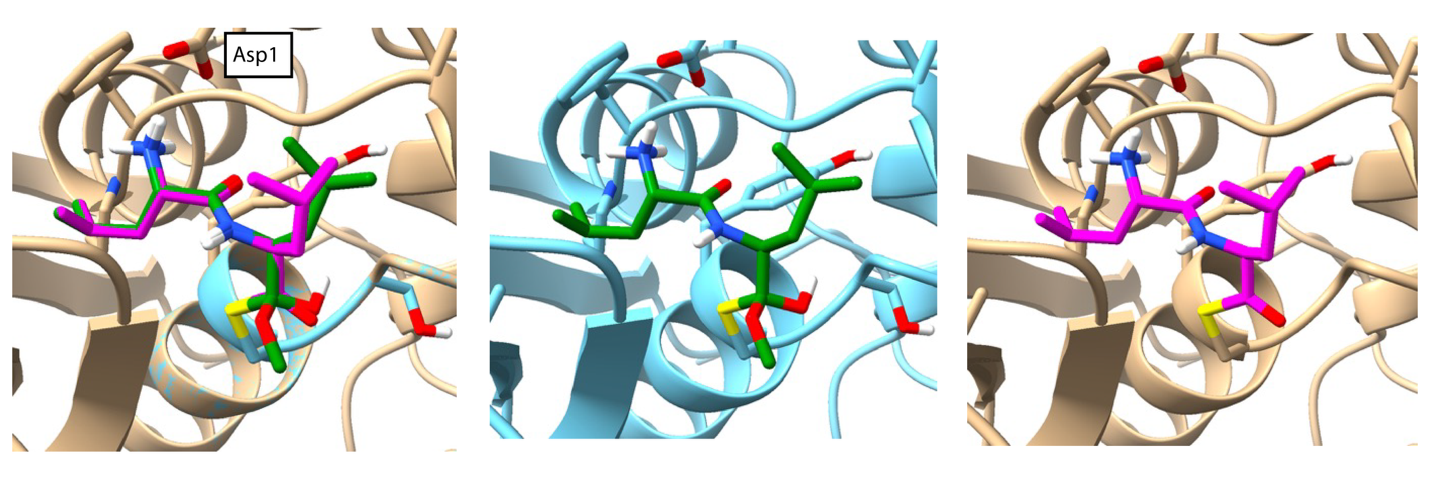

Supplementary Figure 3. Methoxy loss is generally inconsequential in covalent LLOMe-CTSC docking studies (related to Extended Data Figure 4A).

(Left) overlay of docking structures of produced starting from LLOMe in Maestro covalent docking function default mode which retains the methoxy group of the dipeptide after Cathepsin C C234 nucleophilic attack (center, green) vs a custom script which forces the software to produce a thioester and lose the methoxy group (right, magenta). As the structures overlap almost perfectly in the region of interest (amino-terminus of dipeptide) we used the default mode for simplicity and manually deleted the methoxy group to avoid confusion in Extended Data Figure 4A.

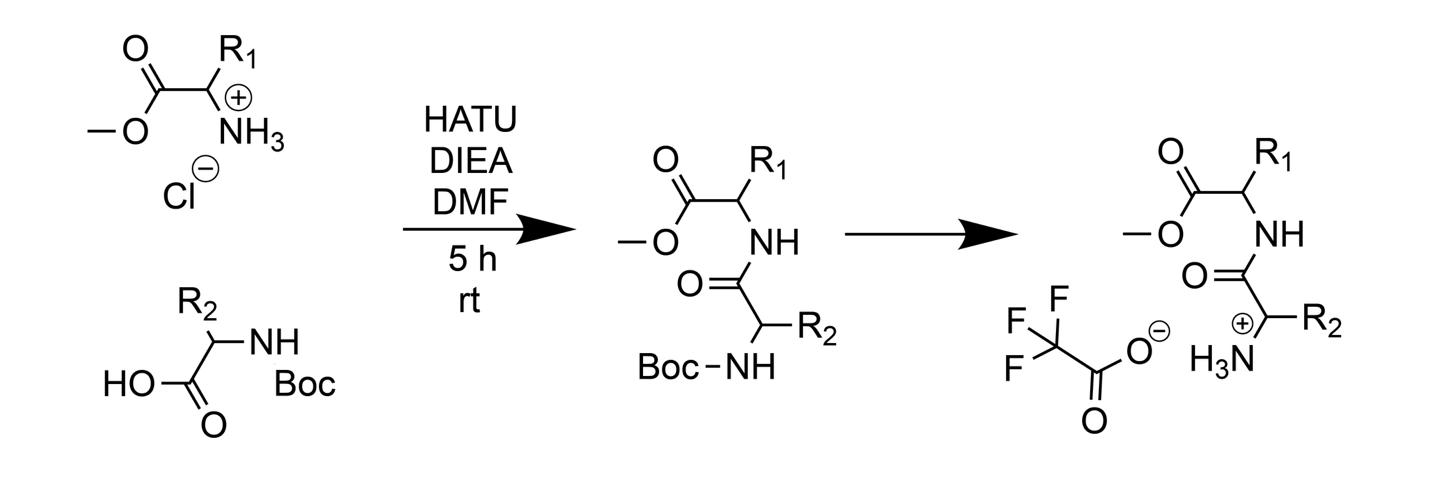

**Supplementary Figure 4. General reaction scheme for the synthesis of dipeptide methyl esters.**

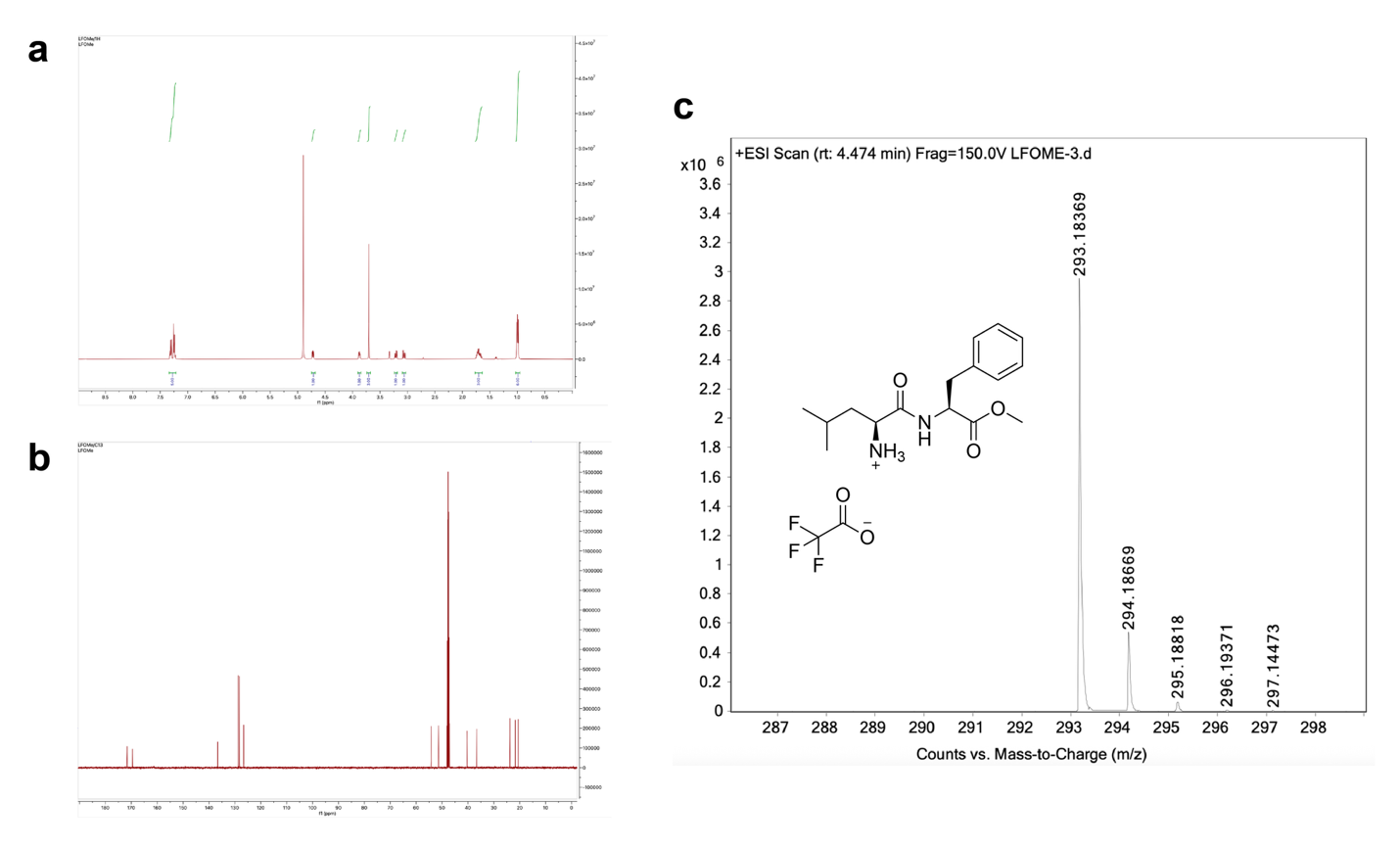

Supplementary Figure 5. Characterization of synthesized dipeptide ester LFOMe.

(**A**) ^1^H NMR (500 MHz, CD_3_OD): 7.29 (2H, m), 7.23 (3H, m), 4.70 (1H, m), 3.86 (1H, m), 3.68 (3H, s), 3.19 (1H, dd), 3.04 (1H, m), 1.74-1.62 (3H, m), 0.99 (3H, d), 0.96 (3H, d). (**B**) ^13^C NMR (500 MHz, CD_3_OD): 173.0, 170.9, 138.0, 130.1, 129.6, 128.0, 55.6, 52.8, 52.7, 41.7, 38.0, 25.2, 23.1, 21.9 (**C**) HRMS-ESI: m/z 293.1837 found, 293.1865 calculated for [M+H] C_16_H_25_N_2_O_3_.

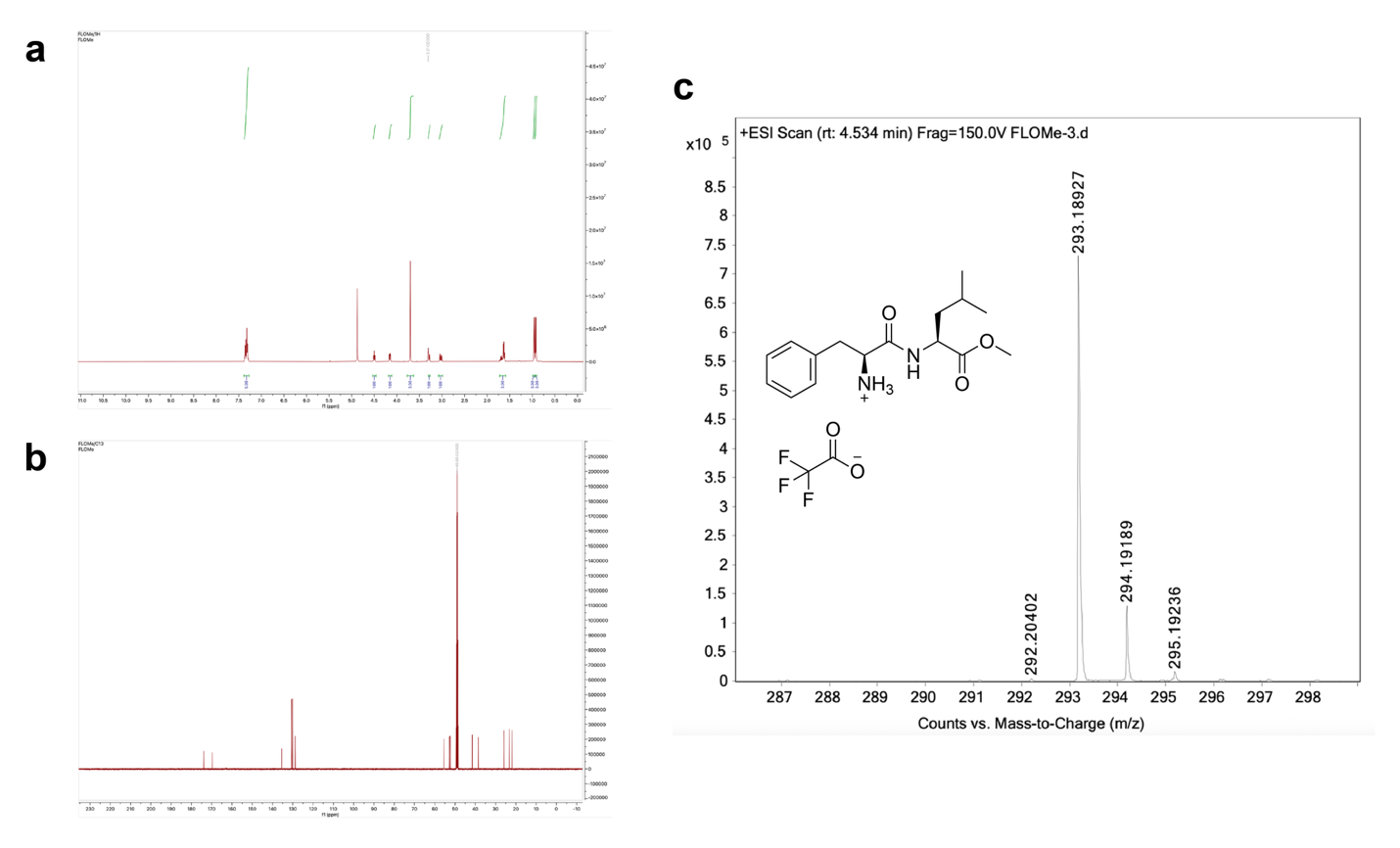

Supplementary Figure 6. Characterization of synthesized dipeptide ester FLOMe.

(**A**) ^1^H NMR (500 MHz, CD_3_OD): 7.33 (5H, m), 4.50 (1H, t), 4.16 (1H, t), 3.70 (3H, s), 3.27 (1H, d), 3.03 (1H, m), 1.59-1.73 (3H, m), 0.96 (3H, d), 0.92 (3H, d). (**B**) ^13^C NMR (500 MHz, CD_3_OD): 173.9, 169.8, 135.5, 130.6, 130.1, 128.8, 55.5, 52.8, 52.3, 41.8, 38.5, 25.8, 23.2, 21.8. (**C**) HRMS-ESI: m/z 293.1893 found, 293.1865 calculated for [M+H] C_16_H_25_N_2_O_3_

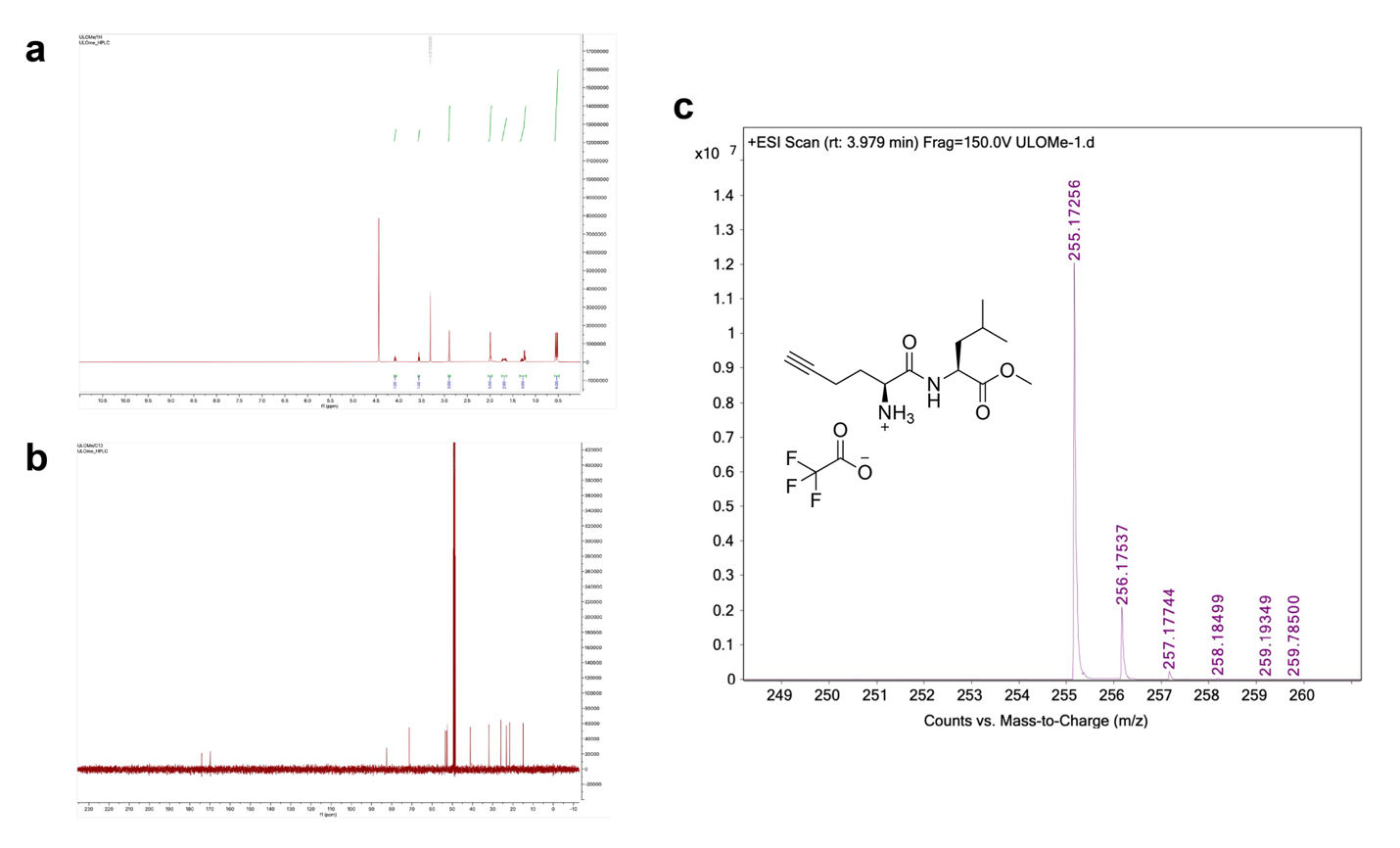
 Supplementary Figure 7. Characterization of synthesized dipeptide ester ULOMe.

(**A**) ^1^H NMR (500 MHz, CD_3_OD): 4.50 (1H, t), 3.98 (1H, t), 3.73 (3H, s), 2.38-2.43 (3H, m), 2.02-2.17 (2H, m), 1.61-1.78 (3H, m), 0.97 (3H, d), 0.94 (3H, d). (**B**) ^13^C NMR (500 MHz, CD_3_OD): 174.1, 169.8, 82.5, 71.3, 53.5, 52.8, 52.4, 41.1, 31.8, 25.9, 23.2, 21.7, 14.9. (**C**) HRMS-ESI: m/z 255.1725 found, 255.1708 calculated for [M+H] C_13_H_22_N_2_O_3_.

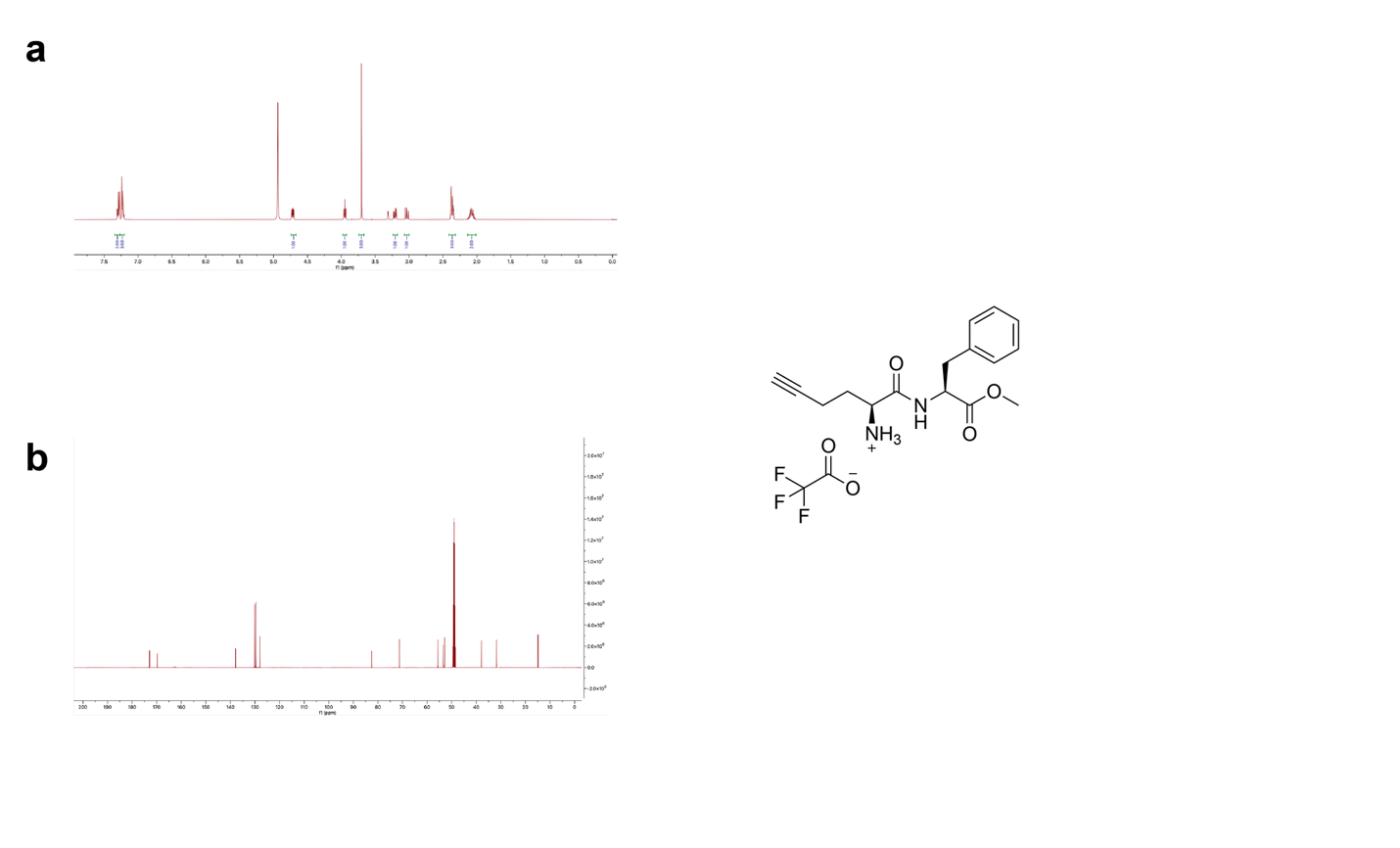
 Supplementary Figure 8. Characterization of synthesized dipeptide ester UFOMe.

(**A**) ^1^H NMR (500 MHz, CD3OD): 7.29 (2H, m), 7.22 (3H, m), 4.71 (1H, m), 3.94 (1H, t), 3.70 (3H, s), 3.20 (1H, dd), 3.02 (1H, dd), 2.36 (3H, m), 2.076 (2H, m). (**B**) ^13^C NMR (500 MHz, CD_3_OD): 172.9, 169.7, 137.9, 130.1, 129.6, 128.0, 82.6, 71.3, 55.6, 53.4, 52.9, 37.9, 31.7, 14.9.

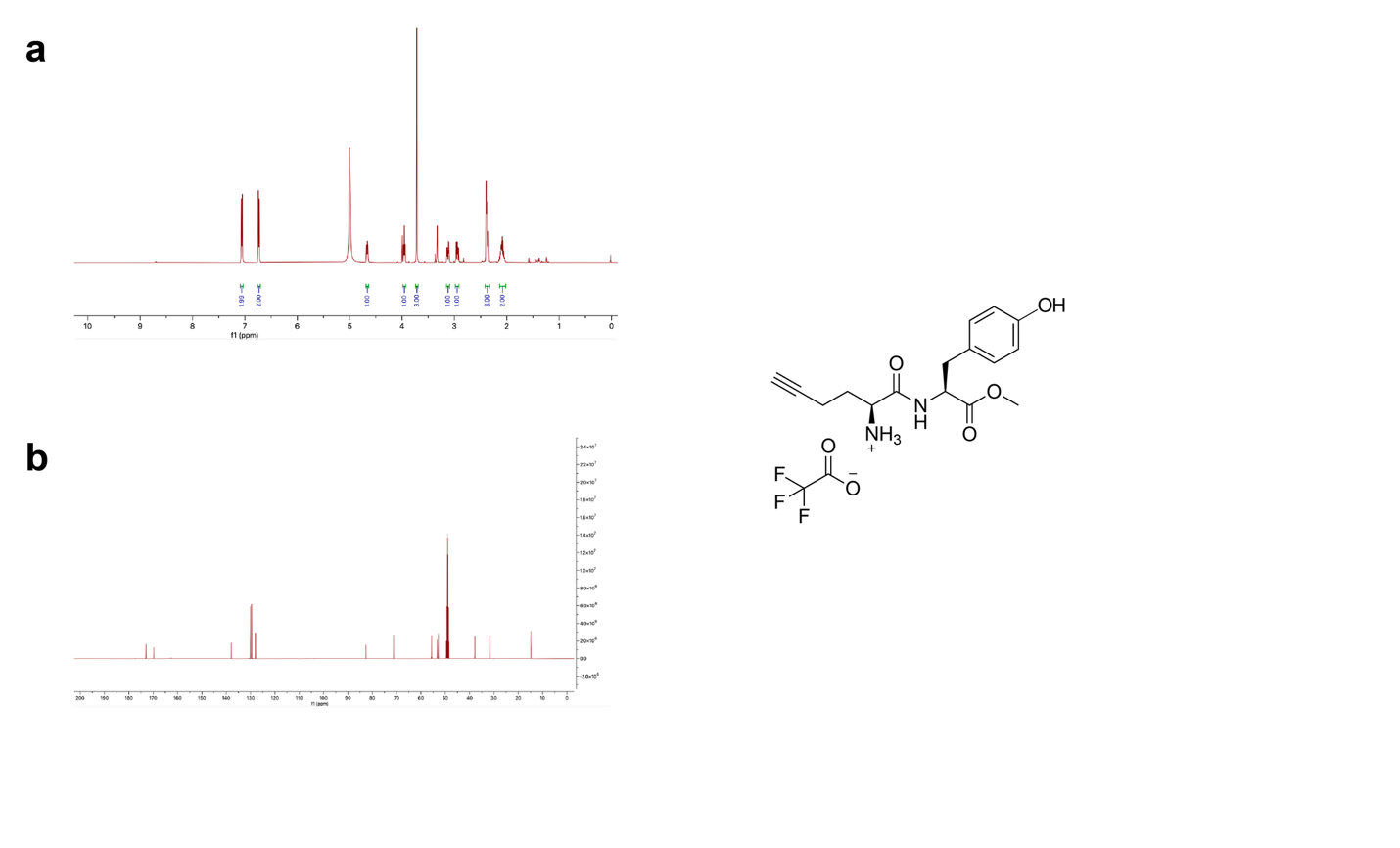
 Supplementary Figure 9. Characterization of synthesized dipeptide ester UYOMe.

(**A**) ^1^H NMR (500 MHz, CD_3_OD): 7.04 (2H, d), 6.71 (2H, d), 4.64 (1H, m), 3.93 (1H, t), 3.10 (1H, dd), 2.92 (1H, m), 2.36 (3H, m), 2.07 (2H, m). (**B**) ^13^C NMR (500 MHz, CD_3_OD): 173.1, 169.7, 167.5, 131.1, 128.5, 116.4, 82.5, 71.3, 55.9, 53.4, 52.8, 37.2, 31.7, 14.9.

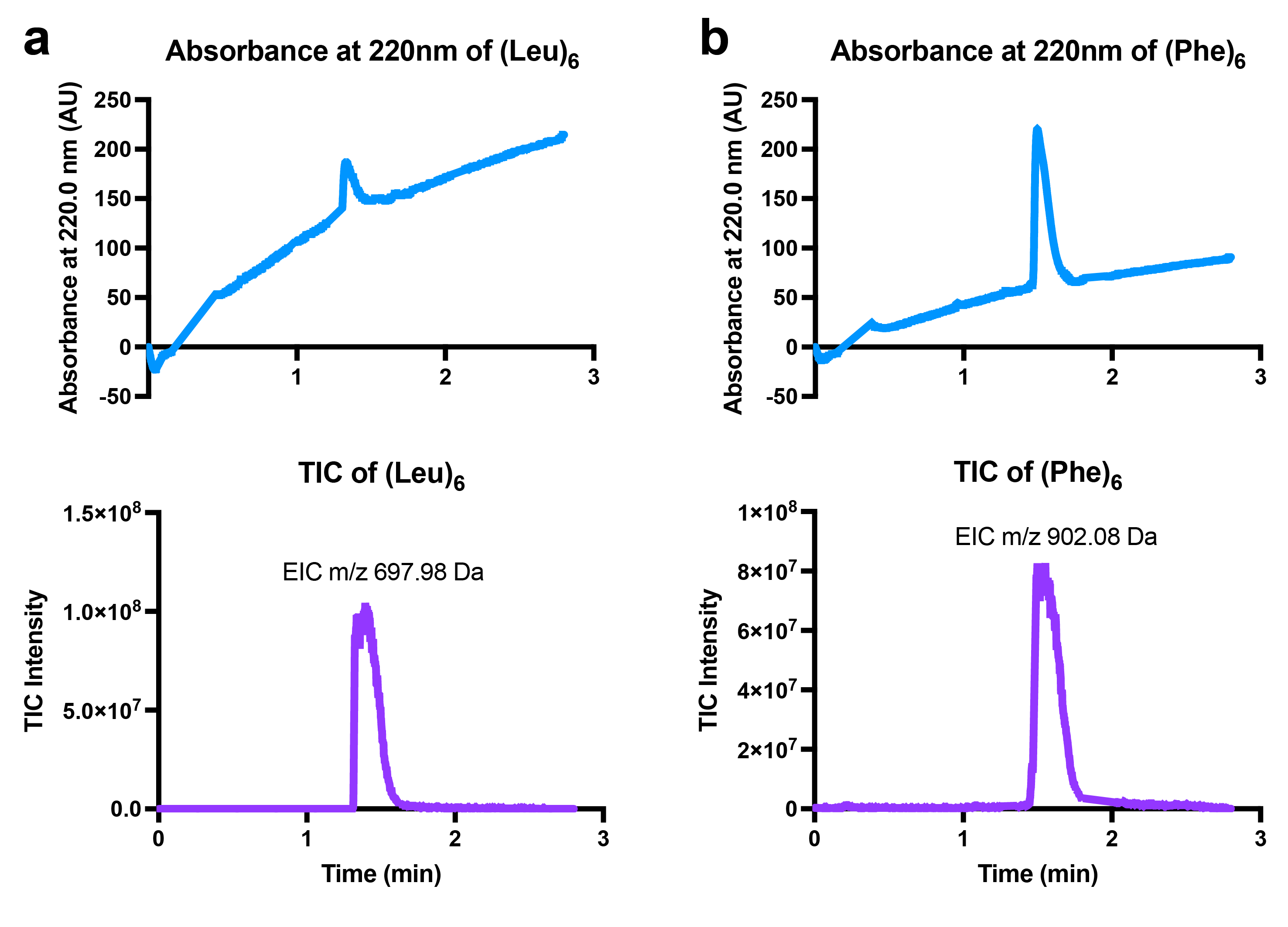
 Supplementary Figure 10. Characterization of synthesized hexapeptides Leu_6_ and Phe_6._

**a**, (Top) LC-MS chromatogram of Leu and (bottom) total ion chromatogram (TIC) of Leu_6_ showing a single dominant peak with correct mass at the expected retention time. **b**, (Top) LC-MS chromatogram of Leu and (bottom) total ion chromatogram (TIC) of Phe_6_ showing a single dominant peak with correct mass at the expected retention time.

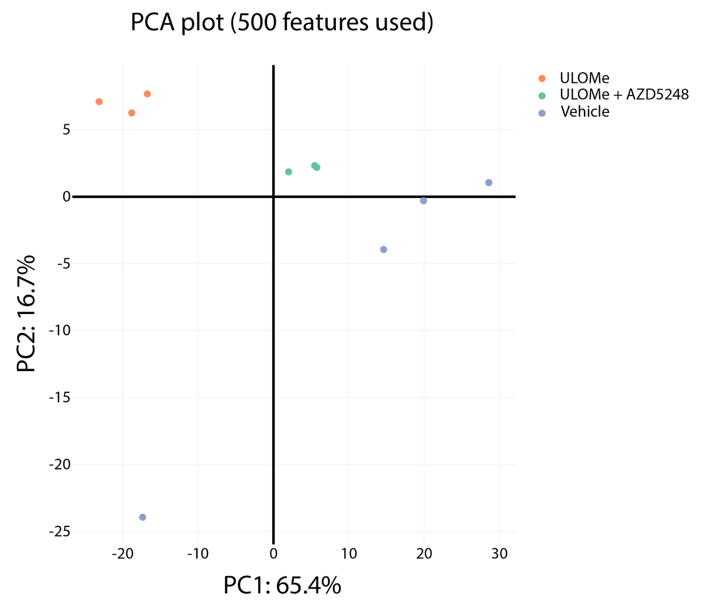

Supplementary Figure 11. PCA plot of ULOMe enriched proteomics dataset.

Principal component analysis (PCA) plot of the ULOMe enriched proteomics dataset displaying clustering of individual ULOMe and ULOMe + AZD5248 samples.

**Supplementary Table 1. ULOMe-based enriched proteomics**

Analysis table of identified protein (650) from the enriched TMT proteomics dataset described in Fig. 4e
