## Extended Data for "Intralysosomal Amyloidogenesis and Proximity Labeling by Cathepsin C"

**Extended Data Figures**

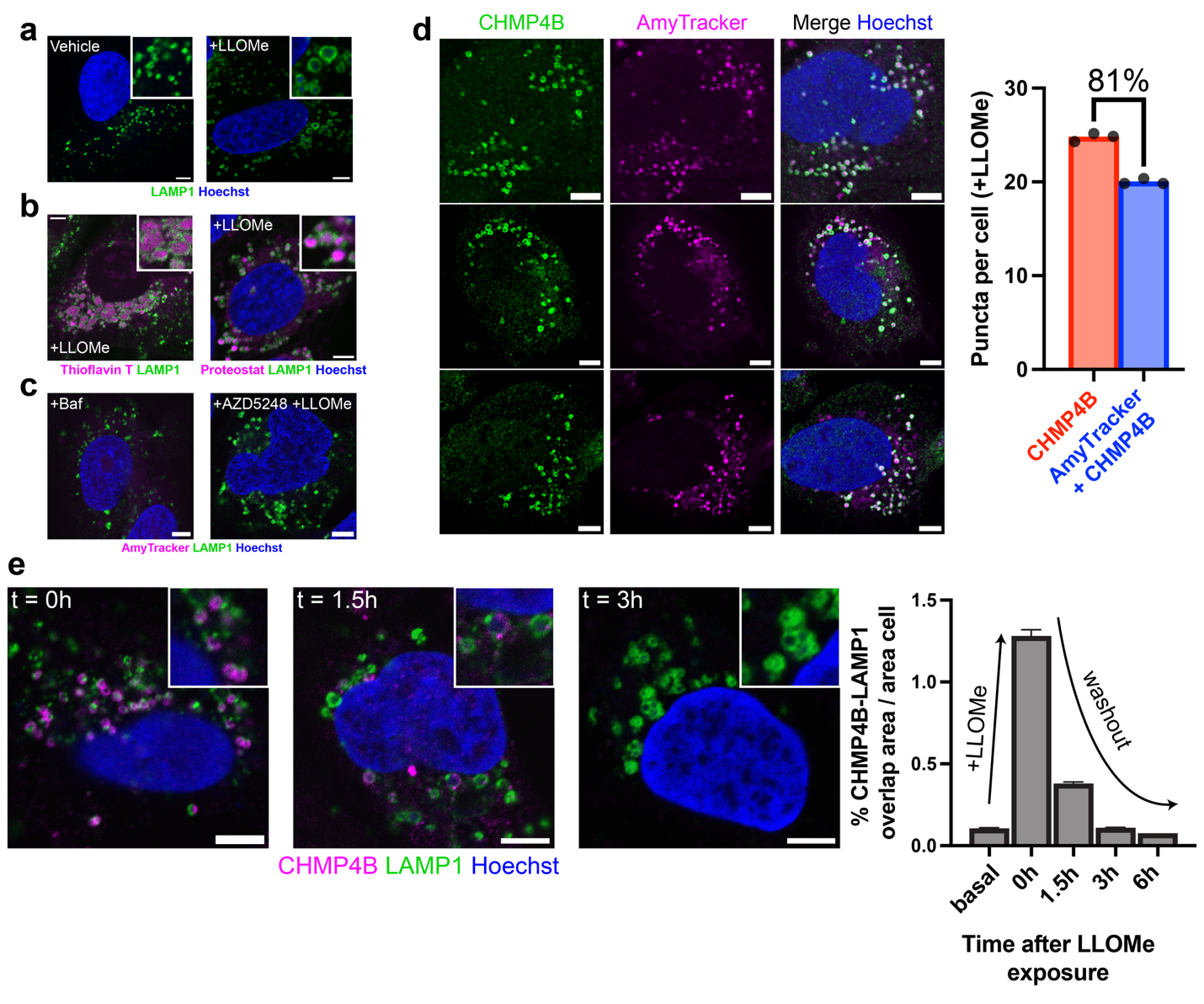
Extended Data Fig. 1. Amyloid deposition by and transient ESCRT response to LLOMe.

Representative images in U-2 OS cells of: a, LAMP1 under basal conditions and upon LLOMe treatment (1 mM, 10 minutes); b, Thioflavin T (1 mM) or (right) Proteostat (2 μg/mL) staining after treatment with LLOMe (1 mM, 10 minutes); c, a lack of AmyTracker staining upon (left) bafilomycin A1 treatment (250 nM, 180 minutes) or (right) pretreatment of CTSC inhibitor AZD5248 (10 μM, 90 minutes) before treatment with LLOMe (1 mM, 60 minutes). d, Additional representative images displaying close association of CHMP4B and AmyTracker puncta after LLOMe treatment (1 mM, 10 minutes) with accompanying comparison of the mean CHMP4B puncta number per cell against AmyTracker-CHMP4B colocalized puncta, the percent mean CHMP4B puncta area associated with AmyTracker is indicated (analysis of n ≥ 145 cells per replicate). e, (Left) representative images of CHMP4B accumulation on LAMP1-positive vesicles at 0, 1.5, and 3-hour timepoints after LLOMe (1 mM, 10 minutes) washout and (right) quantitation of CHMP4B-LAMP1 overlap at the indicated timepoints (analysis of n≥740 cells per timepoint, SEM shown as error bars). Scalebars = 5 μm.

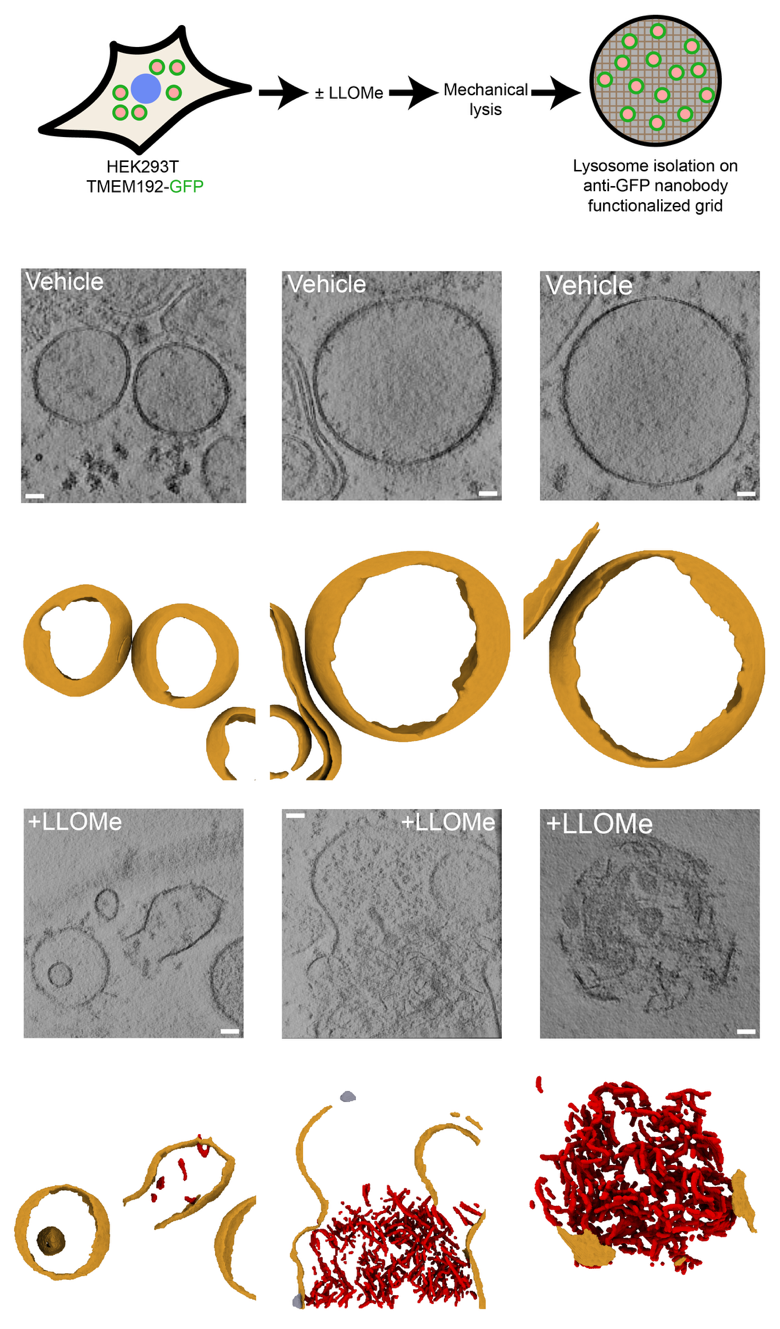

Extended Data Fig. 2. Additional examples of CryoET observation of severe membrane disruption and fibril accumulation in affinity-captured lysosomes.

(Top) Schematic of lysosome affinity-capture strategy from HEK293T for CryoET observation, (bottom) representative tomographic slabs (average of five consecutive slices) and corresponding 3D segmentations of lysosomes from untreated cells (top row) or LLOMe-treated cells (1 mM, 10 min; bottom row). In the segmentations, membranes are colored orange, fibrils red, and flotillin structures gray. Scale bars =10 nm

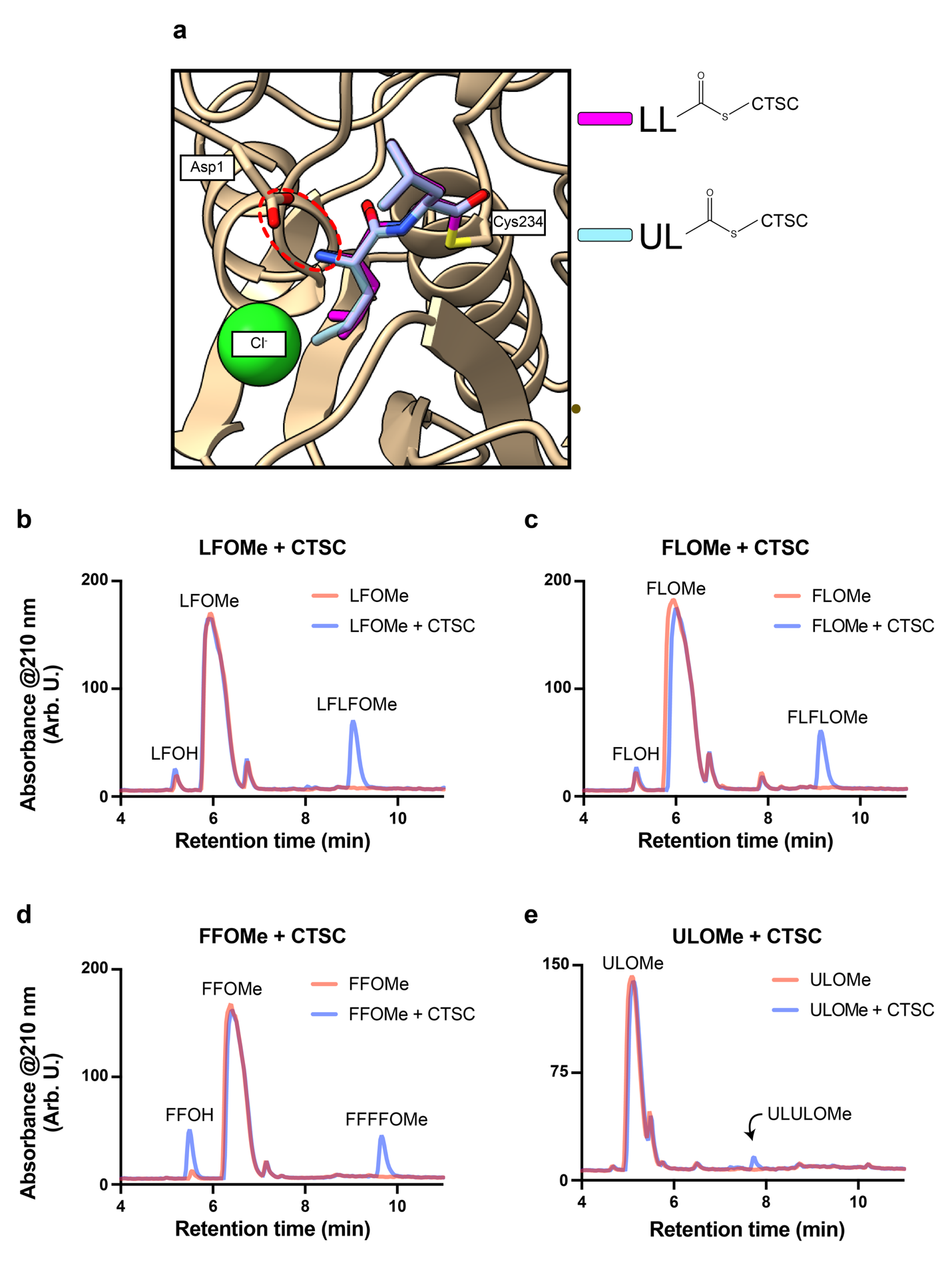
Extended Data Figure 3. Molecular docking of ULOMe and *in vitro* CTSC ligation reactions.

a, Docked poses of CTSC covalently bound to ligands LLOMe (magenta) and ULOMe (light blue) at Cys234 displaying the preservation of the Asp1-dipeptide N-terminus interaction denoted by a dashed red oval. b-e, LC-MS chromatograms of b, LFOMe alone or LFOMe + CTSC; c, FLOMe alone or FLOMe + CTSC; d, FFOMe alone or FFOMe + CTSC; and e, ULOMe alone or ULOMe + CTSC (20 mM dipeptide and 100 nM CTSC respectively, incubated 8 hours at 37 °C in 20 mM NaPi pH 6.5 150 mM NaCl). Labeled peaks were identified by mass.

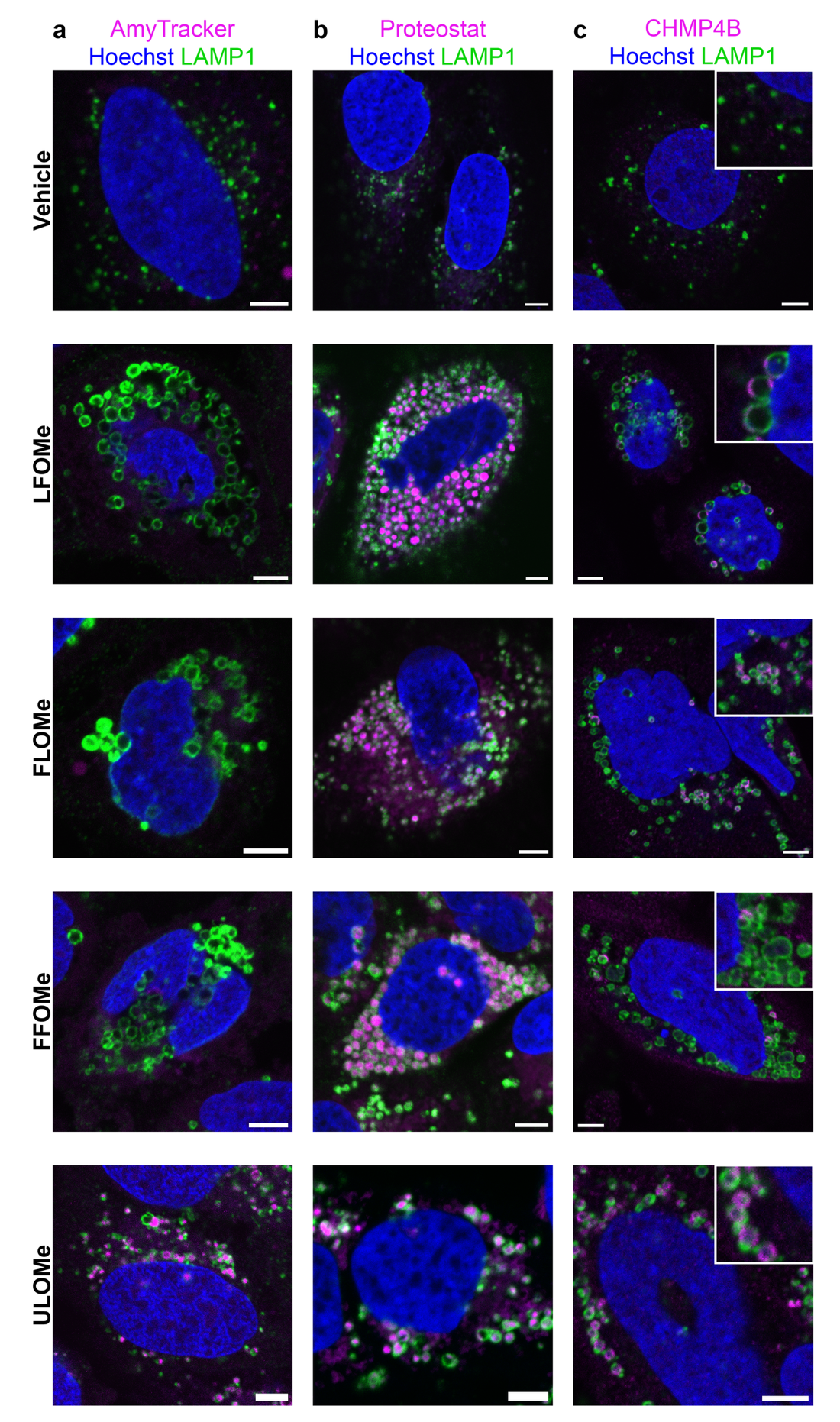

Extended Data Fig. 4. Differential AmyTracker, Proteostat, and CHMP4B staining induced by LLOMe analogs.

Representative images in U-2 OS cells following DMSO treatment, or FL-, FF- or ULOMe treatment (1 mM, 60 minutes) stained against LAMP1 and a, AmyTracker b, Proteostat, or c, CHMP4B. Scalebars = 5 μm.

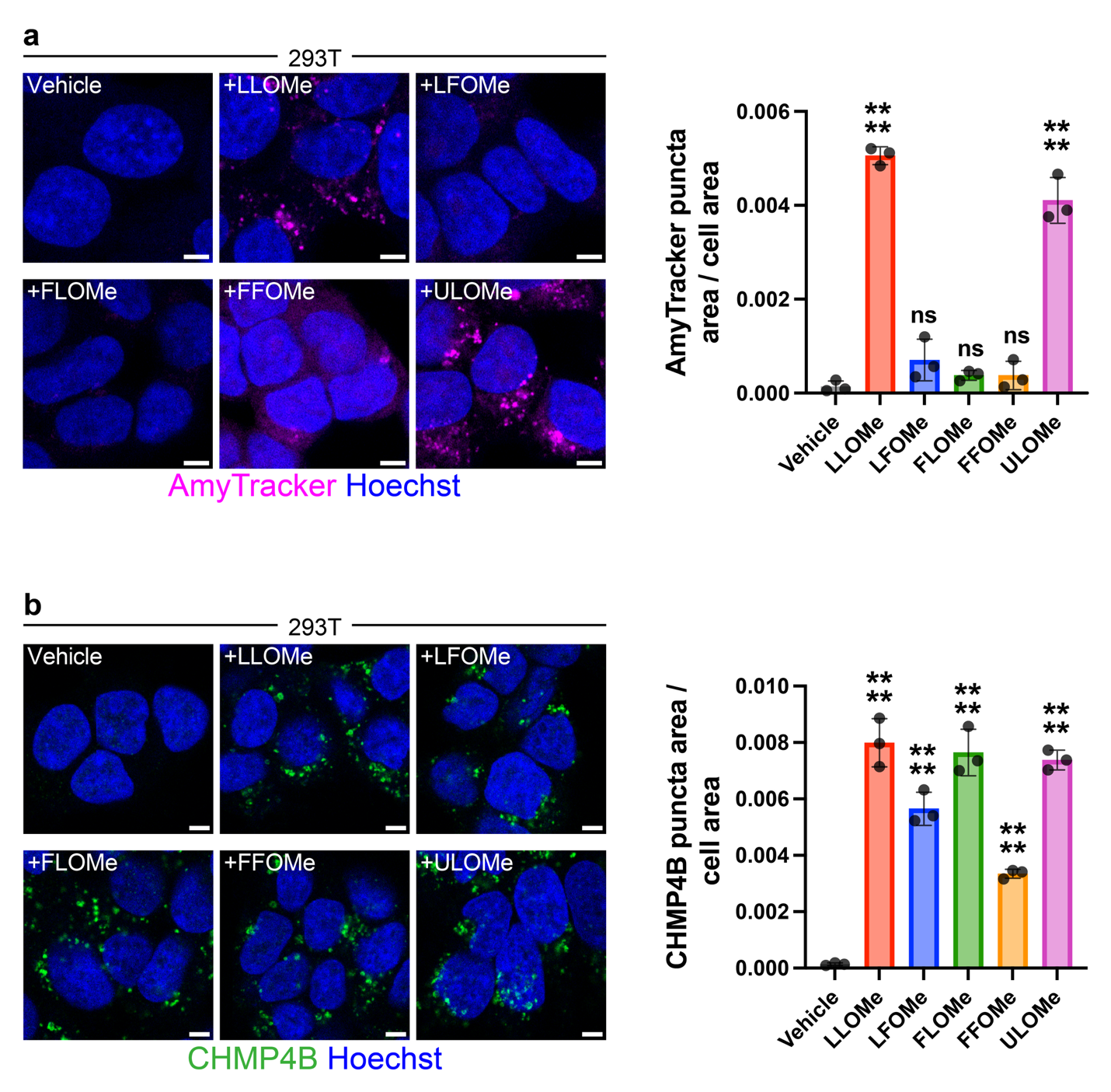
Extended Data Fig 5. Characterization of LLOMe analog phenotypes in HEK293T.

a, (Left) representative images of AmyTracker staining in HEK293T cells treated with DMSO or the indicated dipeptide ester (1 mM, 60 minutes); (right) quantitation of AmyTracker puncta area per cell area per treatment condition (analysis of n ≥ 194 cells per replicate, standard deviation shown as error bars, comparisons against vehicle analyzed using one-way ANOVA, ****=P≤0.0001, ns=P>0.05). b, (Left) representative images of CHMP4B staining in HEK293T cells treated with DMSO or the indicated dipeptide ester (1 mM, 60 minutes); (right) quantitation of CHMP4B puncta area per cell area per treatment condition (analysis of n ≥ 181 cells per replicate, standard deviation shown as error bars, comparisons against vehicle analyzed using one-way ANOVA). Scalebars = 5 μm.

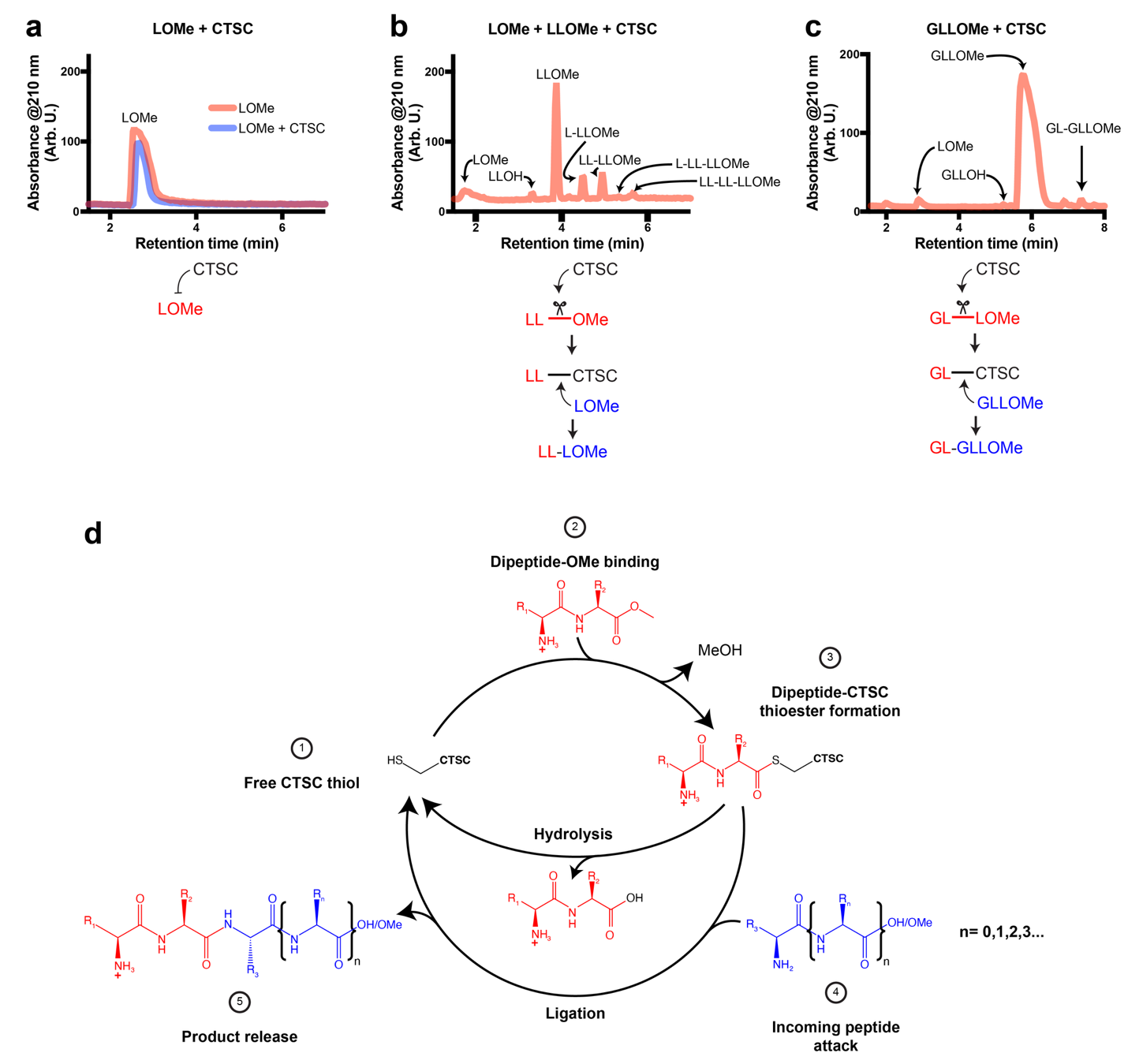
Extended Data Fig. 6. CTSC-catalyzed generation of non-dipeptide repeat products.

LC-MS chromatograms of **a**, LOMe alone or LOMe + CTSC (20 mM and 100 nM respectively); **b**, LOMe + LLOMe + CTSC (10 mM, 10 mM, and 100 nM respectively); and **c**, GLLOMe + CTSC (20 mM and 100 nM respectively) incubated 8 hours at 37 °C in 20 mM NaP_i_ pH 6.5 150 mM NaCl. Labeled peaks were identified by mass. **d**, Schematic of proposed mechanism of CTSC R_1_-R_2_-OMe ligation, where R_1_ and R_2_ belong to the set of natural and unnatural amino acids for which CTSC bears affinity. The formation of a thioester intermediate (3) can resolve through hydrolysis, releasing the dipeptide carboxylic acid, or (4) undergo nucleophilic attack by the N-terminus of an incoming peptide of length n, harboring residues R_3_…R_n_, which may belong to a distinct set of amino acids from R_1_-R_2_. This attack affords the release of a new peptide (5) of length 2+(1+n), composed of R_1_-R_2_-R_3_…R_n_.

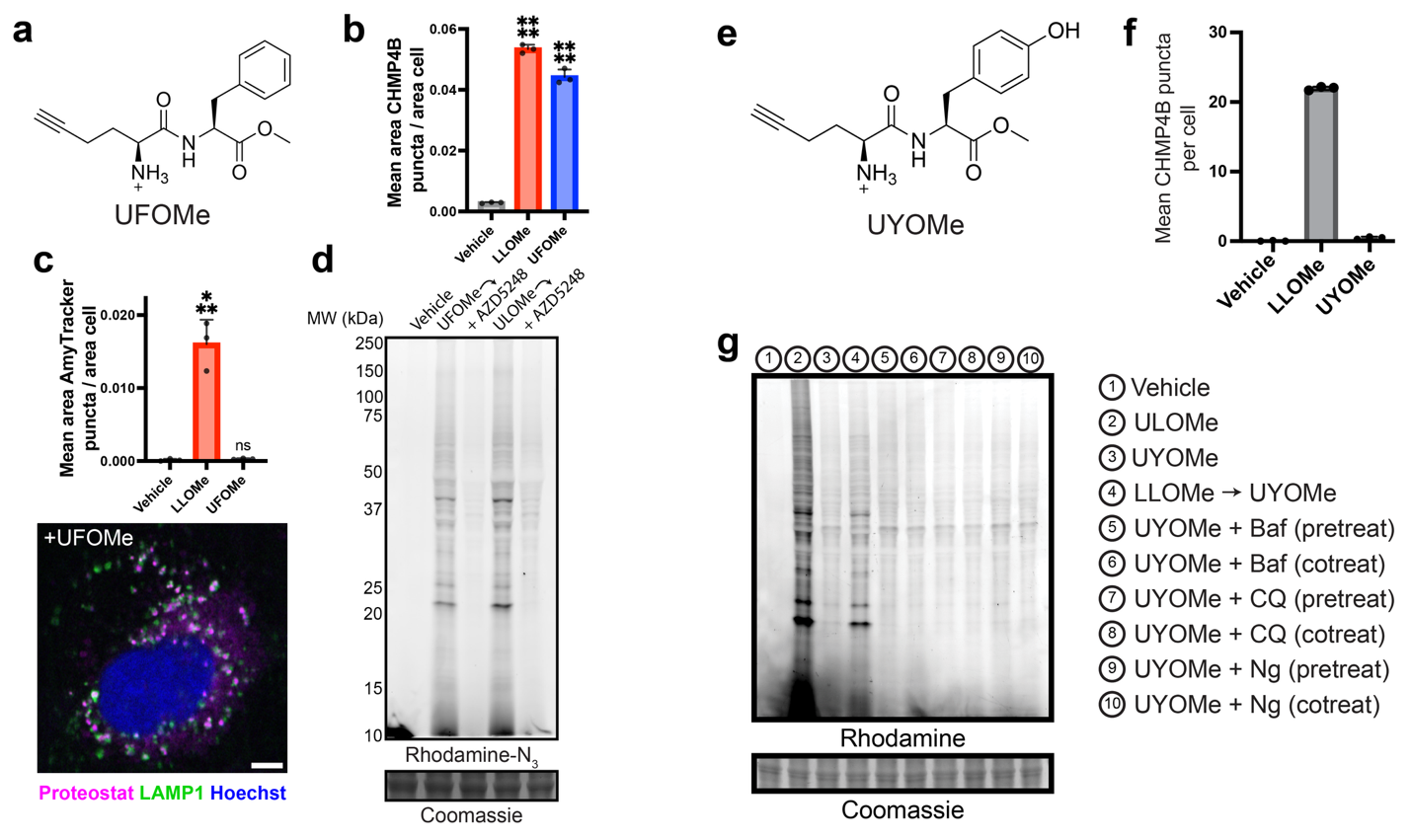
Extended Data Fig. 7. Characterization of UFOMe and UYOMe propensities to induce lysosome damage and LILAC.

**a**, Structure of UFOMe. Comparison of **b**, CHMP4B and **c**, AmyTracker puncta staining in U-2 OS between vehicle, LLOMe (1 mM, 60 minutes), and UFOMe (1 mM, 60 minutes) (analysis of n ≥ 100 cells per treatment, standard deviation shown as error bars, comparisons against vehicle analyzed using one-way ANOVA, ***=P≤0.001, ****=P≤0.0001) and representative image displaying LAMP1-associated Proteostat staining after UFOMe treatment. **d**, In-gel fluorescence displaying LILAC induced by UF- and ULOMe (1 mM, 60 minutes). **e**, Structure of UYOMe. **f**, Quantitation of CHMP4B puncta induced by vehicle, LLOMe, or UYOMe treatment (1 mM, 60 minutes) (analysis of n ≥ 74 cells per treatment). **g**, In-gel fluorescence displaying the lack UY- ligation only upon LLOMe pretreatment (1 mM, 10 minutes LLOMe followed by 1 mM, 60 minutes UYOMe), and not under basal or lysosome deacidifying conditions (250 nM bafilomycin A1, 10 μM chloroquine, or 100 nM nigericin either pretreated alone 90 minutes before 1 mM UYOMe addition, or cotreated with UYOMe). Scalebar = 5 μm.

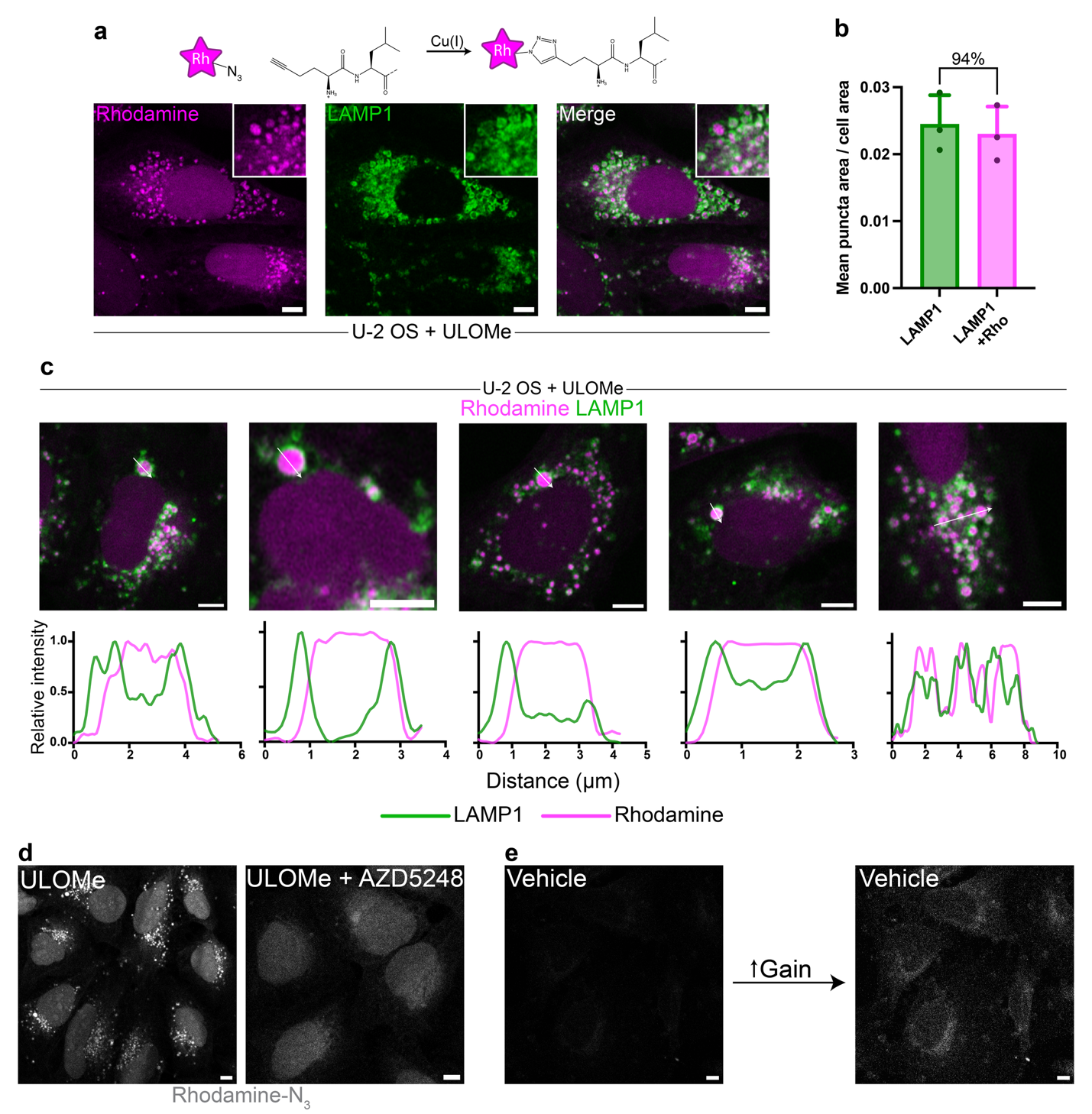
Extended Data Fig. 8. UL-species accumulate in endolysosomes dependent on CTSC activity.

a, (Top) schematic of addition of rhodamine-N_3_ to UL-species by CuAAC, (bottom) representative images displaying endolysosomal and nuclear accumulation of UL-species in U-2 OS cells treated with ULOMe (1 mM, 60 minutes). b, Comparison of the mean LAMP1 puncta number per cell against rhodamine-LAMP1 colocalized puncta; the percent mean LAMP1 puncta area associated with rhodamine is indicated (analysis of n ≥ 86 cells per replicate, standard deviation shown as error bars). c, Line analyses of rhodamine-N_3_ and LAMP1 staining highlighting lumenal accumulation of UL-species. d, Representative images in U-2 OS cells of rhodamine addition by CuAAC after (left) ULOMe treatment (1 mM, 60 minutes) or (right) ULOMe treatment after AZD5248 pretreatment (10 μM, 90 minutes), and e, of rhodamine-N_3_ addition by CuAAC after DMSO treatment. Scalebars = 5 μm.
