## Supplementary material for "Intralysosomal Amyloidogenesis and Proximity Labeling by Cathepsin C": Methods

**Materials and Methods**

Materials

Primary antibodies used were mouse anti-LAMP1 (Cell Signaling catalog no. 15665), rabbit anti-CHMP4B (Proteintech catalog no. 501728932), rabbit anti-LC3B (Cell signaling catalog no. 2775), Mouse anti-Vinculin (Sigma-Aldrich catalog no. V9131), mouse anti-Cathepsin C (Santa Cruz Biotechnology catalog no. sc-74590), and rabbit anti-β-Actin (Cell Signaling catalog no. 4970). Secondary antibodies for imaging, goat anti-mouse Alexa Fluor 488 and goat anti-rabbit Alexa Fluor 647 were purchased from Thermo Fisher Scientific (catalog no. A11001 and A21245 respectively). Secondary Antibodies for immunoblots, goat anti-rabbit-HRP and goat anti-mouse-HRP, were purchased from Cell Signaling (catalog no. 7074 and 7076 respectively) and visualized using SuperSignal West Pico PLUS substrate (Thermo Scientific catalog no. 34580). AmyTracker680 was purchased from Ebba Biotech AB, Proteostat aggresome detection kit was purchased from Enzo (catalog no. ENZ-51035-0025), and thioflavin T was purchased from Sigma-Aldrich (catalog no. 596200). CellTiter Glo assay reagents were purchased from Promega Corporation (catalog no. G9242). Rhodamine-PEG_3_-N_3_ (Supplementary Fig. 2) was synthesized in-house. AZD5248 was purchased from R&D systems inc. (catalog no. 7130), E64d was purchased from Bio-Techne (catalog no. 4545), and bafilomycin A1 was purchased from RPI (catalog no. B40500). Boc-propargyl alanine was purchased from Enamine (EN300-195051). LLOMe and FFOMe were purchased from Cayman Chemical (catalog no. 16008) and Chem-Impex (catalog no. 12097) respectively and characterized by us using ^1^H NMR.

In-vitro CTSC-peptide methyl ester reactions and liquid chromatography-mass spectrometry

For (Leu-Leu)_n_ products, to produce large enough precipitate mass to separate from the reaction mixture, LLOMe solid was dissolved directly into 20 mM sodium phosphate pH 6.5, 250 mM NaCl at a concentration of 100 mM, and the reaction was initiated by addition of 500 nM recombinant mouse active CTSC (R&D Systems catalog no. 2336-CY). The reaction was incubated at 37° C overnight with shaking, and the resulting precipitate was pelleted by centrifugation and air dried overnight after aspiration of supernatant. For LC-MS analysis, pellets were then solubilized in DMSO before injection.

For all other reactions, peptides (from DMSO stocks) and CTSC were mixed at the indicated concentrations under the reaction conditions described above, and an aliquot of the reaction mixture was analyzed by LC-MS after the indicated amount of time.

LC-MS analysis was carried out on an Agilent 1260 Infinity II. Sample was loaded onto a C18 column (Agilent catalog no. 827700-902) and eluted using 2-95% vol/vol acetonitrile gradient supplemented with 0.1% vol/vol TFA.

X-ray diffraction

X-ray diffraction on enzymatically prepared fibrils was carried out on a Bruker Microstar APEX II CCD diffractometer equipped with Cu Kα radiation (λ = 1.54178 Å). LLOMe + CTSC reaction precipitate was mounted on a cryoloop and data was collected at room temperature. A single Phi 360° scan was collected with a sample-to-detector distance of 60 mm and an exposure time of 5 min.

Docking in Maestro Elements software CovDock:

We docked dipeptide methyl esters using the manufacture recommended standard procedure in Maestros Elements (2024-4 release) CovDock software after minimizing the input conformation in ChemDraw pro. A custom docking script was explored but as described in Supplementary Fig. 3, it made little difference to the structure which is dominated by the formation of a salt bride between the protonated dipeptide amino-terminus and the negatively charged Asp 1 residue in the binding site.

General procedure for HATU coupling synthesis of dipeptide methyl esters:

LFOMe, FLOMe, ULOMe, UFOMe, and UYOMe were synthesized on the 100 – 200 mg scale in 40-60% yields using solution phase coupling from their boc- and methyl ester starting amino acids as follows:

General reaction scheme provided in Supplementary Fig. 4. To a solution of the boc-amino acid (1 equiv.) and amino acid-methyl ester HCl salt (1 equiv.) and HATU (1.05 equiv.) in DMF (0.1 M in amino acid substrate) was added DIEA (2 equiv.) before stirring the reaction at rt. After 5 hours, the solvent was removed under reduced pressure and the mixture redissolved in EtOAc (20 mL). The organic phase was washed with brine (3 × 20 mL) and dried over Na_2_SO_4_ before the solvent was removed under reduced pressure. The crude boc-dipeptide methyl ester was purified using flash column chromatography (Teledyne Isco CombiFlash Rf+ system, 4 g SiO_2_, Hexane -EtOAC gradient elution, 10-80% over 14 min.) with products typically eluting first around 40% EtOAc. The purity of this boc protected intermediate was verified by ^1^H NMR before boc cleavage in TFA (1 mL) for 5 min at rt. Unbound TFA was completely removed by repeat co–evaporation with MeOH until the mass remained unchanged (which we verified as sufficient evidence of TFA removal using ^19^F NMR on a representative sample). A small portion was taken for ^1^H, ^13^C and high-resolution mass spec characterization (Supplementary Figs. 5-9)

Solid peptide preparations were stored at -80 °C until used to make up stock solution in DMSO (1 M).

We obtained high resolution masses for all synthesized compounds on an Agilent 6230 ESI-TOF mass spectrometer coupled to an Agilent 1260 liquid chromatography system. All NMR data was obtained on a Bruker Avance Neo 500 MHz spectrometer with a 5mm BBO probe with a Z-axis gradient.

Synthesis of and fibril preparation from hexapeptides

Leu₆ and Phe₆ were synthesized by solid-phase peptide synthesis (SPPS) on Fmoc-Leu-Wang resin and Fmoc-Phe-Wang resin, respectively, employing DIC/Oxyma Pure as coupling reagents at 90°C. Cleavage from the resin was performed using a TFA/TIPS/H₂O cocktail (95:2.5:2.5, v/v/v) for 2 hours at room temperature. The cleaved peptides were precipitated by addition of cold diethyl ether, collected by centrifugation, and washed three times with diethyl ether. Crude products were purified by reverse-phase HPLC and purity confirmed by LC-MS analysis (Supplementary Fig. 10).

To promote a monomeric starting state, lyophilized peptides were dissolved in 1,1,1,3,3,3-hexafluoro-2-propanol (HFIP) at 1 mM and the solvent evaporated under a gentle stream of nitrogen gas yielding peptide films. The films were subsequently resuspended in dimethyl sulfoxide (DMSO) to prepare 1 mM stock solutions, aided by vortexing and bath sonication. Working solutions of 2 µM Leu₆ and Phe₆ were prepared by diluting the DMSO stocks into 50 mM HEPES, 250 mM NaCl buffer (pH 6.5), with a final DMSO concentration of ≤0.2% (v/v). Samples were incubated at room temperature for 4 days without agitation to allow fibril formation. Fibril morphology was subsequently characterized by electron microscopy.

Negative stain transmission electron microscopy (TEM)

For enzymatically prepared samples, carbon-coated copper grids (400 mesh) were glow-discharged and 10 µL of LLOMe + CTSC reaction mixture was adsorbed for 2 minutes. Excess sample was wicked away and grids were negatively stained with 50 µL 2% uranyl formate for 2 minutes. Excess stain was wicked away and the grids were allowed to dry. Samples were analyzed at 80kV with a ThermoFisher Talos L120C transmission electron microscope and images were acquired with a CETA 16M CMOS camera.

For synthetically prepared samples, peptide fibrils were assessed by negative-stain electron microscopy following established protocols^1^. In brief, undiluted fibril suspensions were applied to glow-discharged 400 square Mesh Copper/Rhodium Maxaform grids (Electron Microscopy Sciences), incubated for 1 min and stained with 2 % uranyl formate. Micrographs were collected on a Thermo Fisher Talos F200C transmission electron microscope operating at 200 keV and equipped with a Gatan K2 direct electron detector.

Preparation, data collection and processing for cryo-EM

2 µM sample in 50 mM HEPES, 250 mM NaCl buffer (pH 6.5) was used for cryo-EM. UltraAuFoil 200 mesh R 2/2 grids were glow-discharged under vacuum for 30 s at 15 mA in a Pelco easiGlow 91000 glow discharge cleaning system (Ted Pella). 3.5 µl sample was applied to the front of the grid, incubated for 1 min, and blotted from the back side of the grid with Whatman 1 filter paper for 4-6 s after the liquid spot on the filter paper stopped spreading. Grids were manually plunge-frozen in a 4 °C cold room with >95% humidity.

Cryo-EM datasets were collected on a Talos Arctica TEM (Thermo Fisher) operating at 200 keV. Movies were recorded using a Falcon 4i direct electron detector (Thermo Fisher) at a nominal magnification of 150,000, corresponding to a nominal pixel size of 0.94 Å. Movies were saved in the electron-event representation (EER) format and recorded at a total electron exposure of 50 e− per Å2. All datasets were collected automatically using EPU (v.3.9, Thermo Fisher) with a defocus range of −1.0 to −2.0 μm. EPUs fast exposure navigation was used to collect data at an 8 µm image shift. All datasets were processed using cryoSPARC (v.4.7). A total of 2,626 movies were collected for Leu_6_ and 647 for Phe_6_ and dose-fractioned into 40 frames, motion corrected using patch-based motion correction, followed by patch-based CTF estimation in cryo-SPARC live. Leu_6_ fibrils were traced within cryoSPARC using the template-free mode and a diameter range of 50-100 Å, extracted in a 300 px box, Fourier-cropped to 100 px and subjected to 2D classification. Aligning classes were selected, re-extracted without binning and subjected to three rounds of 2D classification using increasing initial uncertainty factors (2 to 4). Average power spectra were calculated for two representative class averages showing cross-beta separation. Phe_6_ fibrils required picking in crYOLO, following established guidelines^2^. Fibril segment coordinates were imported into cryoSPARC and extracted with a 1000 px box, Fourier-cropped to 100 px and subjected to 2D classification. Aligning classes were selected, re-extracted without binning and subjected to three rounds of 2D classification using increasing initial uncertainty factors (2 to 4). Average power spectra were calculated for two representative class averages showing cross-beta separation.

Cell culture

U-2 OS, HEK293T, and HLF-a cells were acquired from ATCC (HTB-96, CRL-3216, and CCL-199 respectively). HeLa cells, WT and CTSC KO, were acquired from Abcam (AB255928 and AB265822 respectively). GM05659 fibroblasts were acquired from Corielle Institute. Cells were cultured in DMEM (Thermo Fisher Scientific catalog no. 11995073) supplemented with 10% vol/vol fetal bovine serum, 100 U/ml penicillin and 100 μg/ml streptomycin. Cells were maintained at 37° C supplemented with 5% CO_2_.

iPSC culture and microglial differentiation were carried out as previously reported^3^.

Cloning

All constructs were generated using In-Fusion cloning (Takara Bio) according to the manufacturer's specifications. Cathepsin C (CTSC) and catalytically inactive C234A mutant cDNAs were purchased from IDT and cloned into expression vectors containing CAG promoters and ampicillin resistance. The wild-type and C234A constructs were otherwise identical, differing only in the two nucleotide substitutions that abolish catalytic activity at cysteine 234. All plasmids were propagated in One Shot Stbl3 chemically competent E. coli and purified using the ZymoPURE II Plasmid Maxiprep Kit. Plasmid sequences were confirmed by whole-plasmid sequencing (Plasmidsaurus).

Transfection

Transient transfection of CTSC and C234A mutant CTSC constructs was performed in HeLa as previously described^4^. Briefly, PEI-Max was reconstituted to 1 mg/mL in water and used at a 3:1 ratio of PEI to DNA (w/w) per well of a confluent 6-well dish. TagRFP was linked to the C-terminus of CTSC via a P2A self-cleaving peptide sequence, and fluorescence was assessed 48 h post-transfection prior to treatments, after which treatment with ULOMe was conducted as described.

Cryo-electron tomography sample preparation, plunge freezing, data collection, and segmentation

HEK293T cells stably expressing a carboxy-terminal GFP-tagged TMEM192 were treated with 1 mM LLOMe for 10 min. Cell were lysed lysis using a hypotonic homogenization buffer (25 mM Tris-HCl pH 7.5, 50 mM sucrose, 0.2 mM EGTA, 0.5 mM MgCl_2_), followed by application of shearing forces generated using a 23-G syringe. The resulting lysate was immediately mixed with a sucrose buffer on ice (2.5 M sucrose, 0.2 mM EGTA, 0.5 mM MgCl2). The nuclear fraction was separated by centrifugation (1,000xg, 10 minutes, 4ºC). Supernatants were kept at 4ºC before freezing. The TMEM192-GFP positive lysosomes from this lysate were then purified on-grid using electron-microscopy grids functionalized with anti-GFP nanobodies^5,6^.

Plunge freezing was done using a Leica GP2, a Whatman #1 blotting paper and liquid ethane at -180 °C as a cryogen. Lysate was added to the functionalized grids in a series of a total of six 6 µl drops which were blotted away after each addition. Grids were washed with 6 µl PBS and a final 6 µl drop of PBS was added. The functionalized grids were loaded into the chamber, which was set at 4 °C and 75% humidity. Back-side blotting was then done for 5-6 s, and the grid was plunged into liquid ethane and then stored in liquid nitrogen.

All tilt series were collected on one grid on a Krios G4 equipped with an X-FEG electron gun, a Falcon 4i direct electron detector, and the SelectrisX energy filter. The pixel size was set to 1.51 Å per pixel, and the total dose to 62.93 e− Å−2 linearly spread over 31 tilt images spanning a range of –45° to +45° in 3° increments. The software used for data collection was TFS Tomo 5. The movie frames were saved in the EER format.

Motion correction, tilt-series alignment and tomogram reconstruction were all performed using AreTomo3 v1.0.7^7^.

Representative tomograms of untreated and LLOME-treated samples were imported into a local copick^8^ project, and loaded in napari^9^ using the napari-copick plugin. Membranes, flotillin structures, and filamentous structures inside endo-lysosomal vesicles were segmented semi-manually using napari-nnInteractive^10^, and corrected manually using built-in napari segmentation tools. Final visualizations of segmentations were created in UCSF ChimeraX^11^ using the ChimeraX-copick plugin.

Cell viability

Microglia were seeded in a 96-well plate (10,000 cells/well). After 48 hours, cell culture media was replaced with media containing the indicated concentrations of dipeptide. Cell viability was then analyzed using Cell-Titer Glo 2.0 per manufacturer’s instructions. Luminescence data was normalized relative to vehicle control, plotted and fit to a variable slope Hill equation using GraphPad Prism.

Immunostaining

Cells were seeded in 8-well chamber slides (ibidi catalog no. 80807, 30,000 cells/well) and after 24 hours were treated as described. Cells were fixed using 4% wt/vol formaldehyde (Thermo Fisher Scientific catalog no. PI28906) 10 minutes followed by three washes with DPBS (gibco catalog no. 14190-144), permeabilized with 0.5% wt/vol saponin (MilliporeSigma catalog no. 558255) 15 minutes followed by three wases with DPBS and blocked by incubating in 5% wt/vol BSA (Fisher Scientific catalog no. BP1600) at 37° C for 1 hour. Cells were then incubated with primary antibody mixtures (each at a dilution of 1:100) in 5% wt/vol BSA overnight at 4° C. Following three washes with DPBS cells were incubated with corresponding secondary antibody mixtures (each at a dilution of 1:500) in DPBS 1 hour at room temperature. In the case of Amytracker 680 and Proteostat staining, dye (1:500 vol/vol each) was added and incubated alongside secondary antibodies. Samples were then washed three times with DPBS and further stained with Hoechst 33342 (5 μg/mL, Thermo Fisher Scientific, catalog no. 62249) and in a single case thioflavin T (1 mM) for 5 minutes before further washes. Samples were stored in DPBS supplemented with 5 mM NaN_3_ at 4° C before imaging.

Copper-catalyzed Azide-Alkyne Cycloaddition, in-gel fluorescence and western blots

For addition of rhodamine-N_3_ by CuAAC for imaging, cells were seeded as above and treated with ULOMe. After fixation, permeabilization, and blocking cells were incubated in 100 mM Tris pH 7.4, 2% wt/vol BSA, 4 μM rhodamine-N_3_, 1 mM CuSO_4_, 100 μM TBTA, and 5 mM sodium ascorbate for 1 hour. Cells were then washed with DPBS and additionally processed for immunostaining as above.

For addition of rhodamine-N_3_ by CuAAC for in-gel fluorescence, cells were seeded in 6-well plates (300,000 cells/well) and after 24 hours treated with ULOMe or UFOMe as described. Cells were then harvested by trypsinization and cell pellets lysed in 50 μL DPBS containing Roche protease inhibitors by sonication (2 X 10 second pulses, 10% amplitude). Protein concentrations were quantified by DC assay, and samples were normalized to 1.5 mg/mL (25 μL) in the lysis buffer. Samples were treated with click reagent (3 μL) containing TBTA (1.5 μL/sample of 1.7 mM stock in 4:1 DMSO:tBuOH), CuSO4 (0.5 μL/sample of 50 mM stock in H2O), rhodamine-N_3_ (0.5 μL/sample of 1.25 mM stock in DMSO), and freshly prepared TCEP (0.5 μL/sample of 50 mM stock in DPBS). The samples were incubated at room temperature for 1 h. The reaction was quenched with 8.7 μL 4x LDS loading buffer.

Lysates were then resolved on a 14% tris-glycine SDS-PAGE gel (165V, 67 min) and imaged using the rhodamine and Cy5 filter sets. Proteins were transferred to nitrocellulose membrane (semi-wet transfer, 25V, 7 min) following blocking with 5% milk/TBST for 1 h at room temperature. Membranes were then incubated with primary antibody (1:1,000 in 5% milk/TBST) overnight at 4 °C before being washed 3x with TBST and incubated with secondary antibody (1:2,000 in 5% milk/TBST) at room temperature for 1 h. The blots were washed 3x and imaged using the Thermo SuperSignal Pico kit.

Light microscopy and analysis

Stained slides were imaged on a Zeiss LSM 880 laser-scanning confocal microscope with Zen (Black) 2011 SP7 using a Plan-Apo 63x NA 1.4 oil immersion objective, with a pinhole of 90 microns. Hoechst; Ex: 405 nm 1.2%, Em: 410-587 nm, Gain 550. Alexa Fluor 488; Ex: 488 nm 2.0%, pinhole 52 microns, Em: 545-697 nm, Gain 685. Alexa Fluor 555; Ex 514 nm 2.0%, Em: 545-697 nm, Gain 800. AmyTracker 680; Ex: 561 nm 2.0%, pinhole 50 microns, Em: 578-696nm, Gain 625. Alexa Fluor 647; Ex: 633 nm 2.0%, Em: 638-755 nm, Gain 800. Line average 4x with pixel dwell time of 0.55 microseconds.

Additional imaging was performed on a Zeiss Cell Discoverer 7 in confocal mode using a 50x NA 1.2 water immersion objective with a 0.5x tube lens. Hoechst; Ex: 405 nm 3.5%, pinhole 50 microns, Em: 400-595nm, Gain 700. Alexa Fluor 488; Ex: 488 nm 0.6%, pinhole 46 microns, Em: 410-599 nm, Gain 532. Alexa Fluor 555; Ex 514 nm 1.8%, pinhole 39 microns, Em: 400-590, Gain 666. AmyTracker 680; Ex: 561nm 3.0%, pinhole 53 microns, Em: 565-700 nm, Gain 850. Cy5/Alexa Fluor 647; Ex: 633nm 4.0%, pinhole 22 microns, Em: 656-700nm, Gain 625. Line average 4x with a pixel dwell time of 0.42 microseconds.

Image analysis was carried out using Arivis Pro ver. 4.2.0 (Carl Zeiss Microscopy GmbH). Preparation of representative images for publication was performed in ImageJ^12^.

Alkyne enrichment mass spectrometry – treatment and processing

U-2 OS cells were seeded in 15-cm dishes to confluence, one dish per replicate. Cells were treated with either DMSO, ULOMe (1 mM, 60 minutes) or ULOMe after 90-minute preincubation with CTSC inhibitor AZD5248 (10 μM), then washed and harvested by trypsinization, spun at 300 x g 5 minutes, repetitively washed 3 times, then stored at -80 °C until further processing.

ULOMe-treated samples were prepared for TMT10plex mass spectrometry analysis following established protocols^13-15^. Cell pellets were resuspended in 500 μL of cold DPBS and lysed by probe sonication (2 × 15 pulses, 10% power output). Total protein concentration was determined by Pierce BCA assay, and lysates were normalized to 2 mg/mL in 500 μL (1 mg total proteome per sample). Copper-catalyzed azide–alkyne cycloaddition (CuAAC) was performed by addition of 55 µL of click chemistry master mix (30 µL of 1.7 mM TBTA in 4:1 t-BuOH:DMSO, 10 µL of 50 mM CuSO_4_ in H_2_O, 10 µL of freshly prepared 50 mM TCEP in H_2_O, and 10 µL of 10 mM biotin-PEG4-azide (BroadPharm, cat. no. BP-22119) in DMSO) to each sample. Reactions proceeded for 1 h at room temperature with vortexing every 20 min. Proteins were precipitated by sequential addition of ice-cold methanol (600 μL) and chloroform (200 μL), followed by vortexing and centrifugation at 17,500×g for 10 min at 4 °C. Both phases were aspirated without disturbing the protein disk, which was then resuspended in 500 μL methanol and re-pelleted at 17,500×g for 10 min at 4 °C. After complete aspiration of methanol, protein pellets were stored at −80 °C. Pellets were solubilized in 500 μL of 8 M urea in DPBS supplemented with 10 μL of 10% SDS and sonicated to homogeneity. Reduction was performed with 25 μL of 200 mM dithiothreitol (DTT) at 65 °C for 15 min, followed by alkylation with 25 μL of 400 mM iodoacetamide at 37 °C for 30 min. Samples were transferred to 15 mL conical tubes pre-loaded with 4 mL DPBS and 100 µL of 10% SDS, and the sample tube was rinsed with 1 mL DPBS which was then transferred to the conical. Streptavidin agarose beads (Thermo Scientific, cat. no. 20353; 200 µL of washed bead slurry per sample) were added and probe-labeled proteins were enriched for 1.5 h at room temperature with end-over-end rotation. Beads were collected by centrifugation (3,000×g, 2 min) and washed sequentially with 0.2% SDS in DPBS (2 × 10 mL) and DPBS (1 × 5 mL) in the conical tubes; beads were then resuspended and transferred to Eppendorf Protein LoBind tubes, followed by additional washes with HPLC-grade water (2 × 1 mL) and 200 mM EPPS (1 × 1 mL). On-bead digestion was carried out overnight at 37 °C in 200 µL of trypsin solution (2 M urea, 1 mM CaCl2, 10 µg mL-1 trypsin, 200 mM EPPS, pH 8.15). Beads were pelleted at 3,000×g and the supernatant was collected and diluted with 105 µL of acetonitrile (30% final), followed by 6 μL of the appropriate TMT10plex reagent (20 mg/mL in dry acetonitrile). Labeling proceeded for 1.5 h at room temperature with periodic vortexing, then was quenched with hydroxylamine (6 μL of 5% aqueous solution) for 15 min. Samples were acidified with 5 µL of 100% formic acid, combined, and concentrated to dryness by SpeedVac. Samples were desalted using a Sep-Pak C18 cartridge, dried by SpeedVac, and subjected to high-pH reverse-phase fractionation using peptide desalting spin columns (Thermo Scientific, cat. no. 89852), as previously described^13,15^. Desalted peptides were resuspended in 300 μL of 5% acetonitrile/0.1% formic acid (buffer A) by water bath sonication, loaded onto peptide desalting spin columns, and washed twice with water then once with 5% acetonitrile in 10 mM NH_4_HCO_3_. Peptides were eluted in a stepwise acetonitrile gradient into 15 fractions, every fifth of which was pooled (e.g., fractions 1, 6, 11) to yield five combined fractions, then dried by SpeedVac. Each fraction was resuspended in buffer A prior to LC-MS/MS analysis.

Alkyne enrichment mass spectrometry – TMT LC-MS analysis

TMT LC–MS/MS analysis was performed as previously described^13,15^. Fractions were resuspended in buffer A (5% acetonitrile, 0.1% formic acid in water) and analyzed by TMT LC–MS/MS on an Orbitrap Fusion Tribrid mass spectrometer (Thermo Scientific) coupled to an UltiMate 3000 Series Rapid Separation LC system and autosampler (Thermo Scientific Dionex). Peptides were loaded onto an Acclaim PepMap 100 trap column (Thermo Scientific, cat. no. 164535) and separated on either a 75 μm inner-diameter fused-silica capillary column packed with 1.7 μm C18 (Waters Acquity BEH C18, 25 cm) or an EASY-Spray HPLC column (Thermo Scientific, cat. nos. ES902/ES903) at a flow rate of 0.25 μL min-1. Peptides were resolved over a gradient consisting of 10 min at 5% acetonitrile, 150 min from 5–20%, 20 min from 20–45%, and 5 min from 45–95% acetonitrile (all in 0.1% formic acid/H₂O with 0.1% formic acid), followed by column re-equilibration. Data were acquired using an MS3-based TMT method. MS1 survey scans were collected in the Orbitrap (resolution: 120,000; scan range: 400–1,700 m/z; RF lens: 60%; maximum injection time: 50 ms) with dynamic exclusion enabled (repeat count: 1, duration: 15 s). The most abundant precursors per cycle were selected for MS2 and MS3 analysis. MS2 fragmentation used quadrupole isolation (window: 0.7 Th) followed by collision-induced dissociation in the ion trap (normalized collision energy: 35%; maximum injection time: 120 ms). Following each MS2 acquisition, synchronous precursor selection (SPS) was used to isolate up to ten MS2 fragment ions for MS3 analysis. MS3 precursors were fragmented by HCD and detected in the Orbitrap (collision energy: 55; maximum injection time: 120 ms; resolution: 50,000). Charge-state-dependent isolation windows were applied for MS3 precursor selection: for z = 2, the isolation window was set to 1.2; for z = 3–6, the isolation window was set to 0.7.

Alkyne enrichment mass spectrometry – Data processing and analysis

Raw files were processed using FragPipe (v22.0) with MSFragger using the TMT10-MS3 workflow^16,17^. Spectra were searched against a reviewed human UniProt database (UniProtKB/Swiss-Prot, UP000005640, release 2025-02-12) appended with decoy sequences (“rev_” prefix) and common contaminants. Trypsin (strict, C-terminal cleavage at K/R) was specified as the digestion enzyme with up to two missed cleavages permitted. Peptide length was restricted to 7–50 amino acids with a digest mass range of 200–5,000 Da. Precursor mass tolerance was ±20 ppm and fragment mass tolerance was ±0.6 Da; isotope errors of 0/1/2/3 were considered. Static modifications were applied for cysteine carbamidomethylation (+57.021 Da), TMT10plex on lysine (+229.163 Da), and TMT10plex on peptide N-termini (+229.163 Da). Variable modifications included methionine oxidation (+15.995 Da) and N-terminal protein acetylation (+42.011 Da). The TMT reporter ion m/z region (125.5–131.5 Th) was excluded from fragment matching. PSM validation was performed using Percolator with target-decoy competition post-processing. Protein inference and FDR estimation were performed with ProteinProphet and Philosopher, applying sequential and picked-protein FDR at a 1% threshold at the protein level. TMT10plex reporter ion intensities were extracted and quantified at the MS3 level using TMT-Integrator, with quantification grouped by gene.6 PSMs with peptide probability below 0.9, isolation purity below 0.5, or a reporter ion intensity contribution below 5% of the total signal were excluded. No PSM-level or protein-level normalization was applied between channels. Protein intensities were log_2_-transformed prior to downstream analysis.

Downstream statistical analysis was performed using FragPipe Analyst^18^. Gene-level quantification tables output by TMT-Integrator were imported without additional normalization. Proteins were retained for analysis if they had at least two valid quantification values within at least one experimental condition; all others were excluded. Differential abundance analysis was performed using the limma empirical Bayes framework, with pairwise comparisons conducted between all conditions. P-values were adjusted for multiple comparisons using the Benjamini–Hochberg method. Proteins were considered differentially abundant if they met both an adjusted p-value threshold of 0.05 and an absolute log2 fold-change threshold of 1. Sample clustering and outliers were evaluated using Principal component analysis (Supplementary Fig. 11).

**References**

1	Chowdhury, S., Ketcham, S. A., Schroer, T. A. & Lander, G. C. Structural organization of the dynein–dynactin complex bound to microtubules. *Nature Structural & Molecular Biology* **22**, 345-347 (2015). <https://doi.org/10.1038/nsmb.2996>

2	Wagner, T. *et al.* SPHIRE-crYOLO is a fast and accurate fully automated particle picker for cryo-EM. *Communications Biology* **2**, 218 (2019). <https://doi.org/10.1038/s42003-019-0437-z>

3	Trudler, D. *et al.* Soluble α-synuclein–antibody complexes activate the NLRP3 inflammasome in hiPSC-derived microglia. *Proceedings of the National Academy of Sciences* **118**, e2025847118 (2021). <https://doi.org/doi:10.1073/pnas.2025847118>

4	Longo, P. A., Kavran, J. M., Kim, M. S. & Leahy, D. J. Transient mammalian cell transfection with polyethylenimine (PEI). *Methods Enzymol* **529**, 227-240 (2013). <https://doi.org/10.1016/b978-0-12-418687-3.00018-5>

5	Wang, F. *et al.* General and robust covalently linked graphene oxide affinity grids for high-resolution cryo-EM. *Proc Natl Acad Sci U S A* **117**, 24269-24273 (2020). <https://doi.org/10.1073/pnas.2009707117>

6	Wang, F. *et al.* Amino and PEG-amino graphene oxide grids enrich and protect samples for high-resolution single particle cryo-electron microscopy. *J Struct Biol* **209**, 107437 (2020). <https://doi.org/10.1016/j.jsb.2019.107437>

7	Peck, A. *et al.* AreTomoLive: automated reconstruction of comprehensively corrected and denoised cryo-electron tomograms in real time and at high throughput. *Nature Methods* (2026). <https://doi.org/10.1038/s41592-026-03093-y>

8	Ermel, U. H. *et al.* copick: An open dataset interface and toolkit for collaborative annotation and analysis of cryo-electron tomography data. *Protein Science* **35**, e70578 (2026). <https://doi.org/https://doi.org/10.1002/pro.70578>

9	Ahlers, J. *et al.* napari: a multi-dimensional image viewer for Python. *Zenodo* (2023).

10	Isensee, F. *et al.* nninteractive: Redefining 3d promptable segmentation. *arXiv preprint arXiv:2503.08373* (2025).

11	Goddard, T. D. *et al.* UCSF ChimeraX: Meeting modern challenges in visualization and analysis. *Protein Sci* **27**, 14-25 (2018). <https://doi.org/10.1002/pro.3235>

12	Schindelin, J. *et al.* Fiji: an open-source platform for biological-image analysis. *Nature Methods* **9**, 676-682 (2012). <https://doi.org/10.1038/nmeth.2019>

13	Goetzke, F. W. *et al.* Complexoform-restricted covalent TRMT112 ligands that allosterically agonize METTL5. *Nature Chemical Biology* **22**, 770-782 (2026). <https://doi.org/10.1038/s41589-025-02099-5>

14	Hayward, R. E. *et al.* Tryptoline Stereoprobe Elaboration Identifies Inhibitors of the GRPEL1-HSPA9 Chaperone Complex. *bioRxiv*, 2025.2010.2020.683548 (2025). <https://doi.org/10.1101/2025.10.20.683548>

15	Njomen, E. *et al.* Multi-tiered chemical proteomic maps of tryptoline acrylamide–protein interactions in cancer cells. *Nature Chemistry* **16**, 1592-1604 (2024). <https://doi.org/10.1038/s41557-024-01601-1>

16	Kong, A. T., Leprevost, F. V., Avtonomov, D. M., Mellacheruvu, D. & Nesvizhskii, A. I. MSFragger: ultrafast and comprehensive peptide identification in mass spectrometry–based proteomics. *Nature Methods* **14**, 513-520 (2017). <https://doi.org/10.1038/nmeth.4256>

17	Yu, F. *et al.* Analysis of DIA proteomics data using MSFragger-DIA and FragPipe computational platform. *Nature Communications* **14**, 4154 (2023). <https://doi.org/10.1038/s41467-023-39869-5>

18	Hsiao, Y. *et al.* Analysis and Visualization of Quantitative Proteomics Data Using FragPipe-Analyst. *Journal of Proteome Research* **23**, 4303-4315 (2024). <https://doi.org/10.1021/acs.jproteome.4c00294>
